# IPANEMAP: Integrative Probing Analysis of Nucleic Acids Empowered by Multiple Accessibility Profiles

**DOI:** 10.1101/2020.07.03.186312

**Authors:** Afaf Saaidi, Delphine Allouche, Mireille Regnier, Bruno Sargueil, Yann Ponty

## Abstract

The manual production of reliable RNA structure models from chemical probing experiments benefits from the integration of information derived from multiple protocols and reagents. However, the interpretation of multiple probing profiles remains a complex task, hindering the quality and reproducibility of modeling efforts.

We introduce `IPANEMAP`, the first automated method for the modeling of RNA structure from multiple probing reactivity profiles. Input profiles can result from experiments based on diverse protocols, reagents, or collection of variants, and are jointly analyzed to predict the dominant conformations of an RNA.

`IPANEMAP` combines sampling, clustering, and multi-optimization, to produce secondary structure models that are both stable and well-supported by experimental evidences. The analysis of multiple reactivity profiles, both publicly available and produced in our study, demonstrates the good performances of `IPANEMAP`, even in a mono probing setting. It confirms the potential of integrating multiple sources of probing data, informing the design of informative probing assays.

**Availability:** `IPANEMAP` is freely downloadable at https://github.com/afafbioinfo/IPANEMAP

Supplementary information available at NAR online.

## Introduction

Historically used as a validation assays (Knapp, 1989), enzymatic and chemical probing is increasingly used in combination with computational methods to inform a rational prediction of secondary structure models for RNA (Mathews et al., 2004). Such an integrated approach to structure modeling has led to sizable improvements in prediction accuracy (Washietl et al., 2012) and is currently at the core of successful modeling strategies (Miao et al., 2017). However, the interpretation of probing data, to inform structure prediction, is challenged by a number of factors, including structural heterogeneity, experimental errors, structural dynamics and the potential variability of reactivity measurements across replicates.

Reagents used within selective 2’ -hydroxyl acylation analyzed by primer extension (SHAPE) (Wilkinson et al., 2006) protocols represent a popular class of probes. They react with the 2’-hydroxyl of flexible ribose (McGinnis et al., 2012), although the exact properties observed by SHAPE remain the object of ongoing investigations (McGinnis et al., 2012; Sexton et al., 2017; Hurst et al., 2018; Mlýnský and Bussi, 2018; Frezza et al., 2019; Busan et al., 2019). As ribose flexibility is proportional to the degree of freedom of the nucleotide, it is assumed that SHAPE reagents discriminate nucleotides involved in stable interactions. Different SHAPE reagents are endowed with different dynamics that can help differentiate Watson-Crick base pairs from more dynamic tertiary contacts (Gherghe et al., 2008; Steen et al., 2012; Rice et al., 2014; Busan et al., 2019). Other chemical probes, harbouring different chemical reactivities, have been developed. Amongst the most popular, DiMethyl Sulfate (DMS) is a small molecule that methylates Adenines and Cytosines if not involved in a hydrogen bond, while CMCT reacts with the Watson-Crick face of unpaired Guanosines and Uracils (Ehresmann et al., 1987; Brunel and Romby, 2000). These two reagents not only reveal Watson-Crick base pairing, but also other types of contacts involving the same edge. The diversity of probes, some of which are usable *in vivo* (Zaug and Cech, 1995; Paillart et al., 2004), not only increases coverage of the different positions and structural contexts, but also provides different qualitative information. Modeling can also benefit from a joint analysis of reactivity profiles of single-point mutants, assuming structural homology (Cordero et al., 2012b). Integration of multiple sources of probing to improve structure prediction has thus been widely used since the very early days of RNA structure probing (Moazed et al., 1986; Ehresmann et al., 1987; Romaniuk et al., 1988; Butcher and Burke, 1994; Brunel and Romby, 2000; Cordero and Das, 2015; Somarowthu et al., 2015; Gross et al., 2017) but, to the best of our knowledge, has never been fully automated within a soft constraint framework (Lorenz et al., 2016).

Computationally, the reactivity of a nucleotide is typically used as a proxy to assess the unpaired nature of individual nucleotides. The past couple of decades have nevertheless seen a series of paradigm shifts in the ways probing information is integrated, somehow mirroring the evolution of *ab initio* methods for secondary structure prediction. The seminal work of Mathews (Mathews et al., 2004) used cutoffs to transform reactivity values into hard constraints. Depending on the used reagent/enzyme, significantly unreactive or reactive positions were forced to remain paired or unpaired within predicted models. However, such *hard constraints* can be overly sensitive to the choice of the cutoff, leading to artificially unpaired predictions or unsatisfiable sets of constraints. With the advent of SHAPE probing (Smola et al., 2015), new methods (Deigan et al., 2009; Zarringhalam et al., 2012; Washietl et al., 2012) lifted the requirement of a threshold, supplementing Turner nearest-neighbor energy model (Turner and Mathews, 2010) with pseudo-energies derived from the reactivities. They then performed a (pseudo) energy minimization, optimizing a tradeoff between the thermodynamic stability and its compatibility with the reactivity profile. Popular packages for secondary structure prediction, such as RNAStructure (Mathews et al., 2004) or the Vienna package (Lorenz et al., 2011), now routinely accept SHAPE-like reactivity profiles as input of their main structure prediction methods. Besides free-energy optimization, such methods notably include the joint folding of aligned RNAs (Lavender et al., 2015), partition function, statistical sampling and Maximal Expected Accuracy predictions (Spasic et al., 2017)…

However, independent computational predictions for the same RNA using probing data obtained with different probes often yield substantially different models, none of which are fully consistent with all the probing data. It follows that, while theoretically informative, a multiprobing strategy often leaves a user with different models, from which one cannot objectively decide. In order to identify the best fit with all the data, researchers resort to very intuitive and manual methods such as projecting the results obtained with one probe onto the different models obtained using the constraints from another probe (Herbreteau et al., 2005; James and Sargueil, 2008; Weill et al., 2004). At the end of the modeling process, the modeler is often left with several alternatives, all of which may appear equally consistent with the probing data. Thus, we have developed a new modeling procedure that jointly takes into account multiple probing data, and ultimately yields a small collection of secondary structure models.

## Material and methods

### The IPANEMAP method

We introduce the Integrative Probing Analysis of Nucleic Acids Empowered by Multiple Accessibility Profiles (`IPANEMAP`), a novel approach that integrates the signals produced by various probing experiments to predict one or several secondary structure models for a given RNA. It takes as input one or several reactivity profiles produced in various *conditions*, broadly defined to denote the conjunction of a reagent, a probing technology, ionic concentrations and, in extreme cases, structurally homologous mutants.

It performs a structural clustering across multiple sets of structures sampled in different experimental conditions, and ultimately returns a set of structures representing dominant conformations supported across conditions. Its underlying rationale is that the prominent presence of a stable secondary structure within the (pseudo-)Boltzmann ensembles induced by multiple experimental conditions should increase its likelihood to be (one of) the native structure(s) for a given RNA. It is thus hoped that integrating several reactivity profiles may be used to promote the native structure as one of the dominant structures within the multi-ensemble, and help circumvent the limitation of pseudo-energies derived from single reactivity profile, which are generally not sufficient to elect the native structure as its minimum (pseudo)-free energy candidate. In other words, combining ensembles of structures generated using multiple probing experiments is likely to denoise the Boltzmann (multi-)ensemble, and thus mitigate systematic biases induced by experimental conditions and reagents.

#### Sampling the pseudo-Boltzmann ensembles

Our method, summarized in Figure 1, takes as input a set *𝒟* of probing experiments, each materialized by a reactivity profile. It starts by producing (multi)sets of representative structures for each of the reactivity profiles using a SHAPE-directed variant of the classic Ding-Lawrence algorithm (Ding and Lawrence, 2003). Following Deigan et al. (Deigan et al., 2009), *soft constraints* are used to complement the free energy contributions of the classic Turner energy model with pseudo energy contributions resulting from the reactivity derived from a probing experiment. Given a reactivity *r*_*i*_ for a position *i* in a probing *d*, we associate a free-energy bonus to unpaired positions, defined as

**Figure 1:**
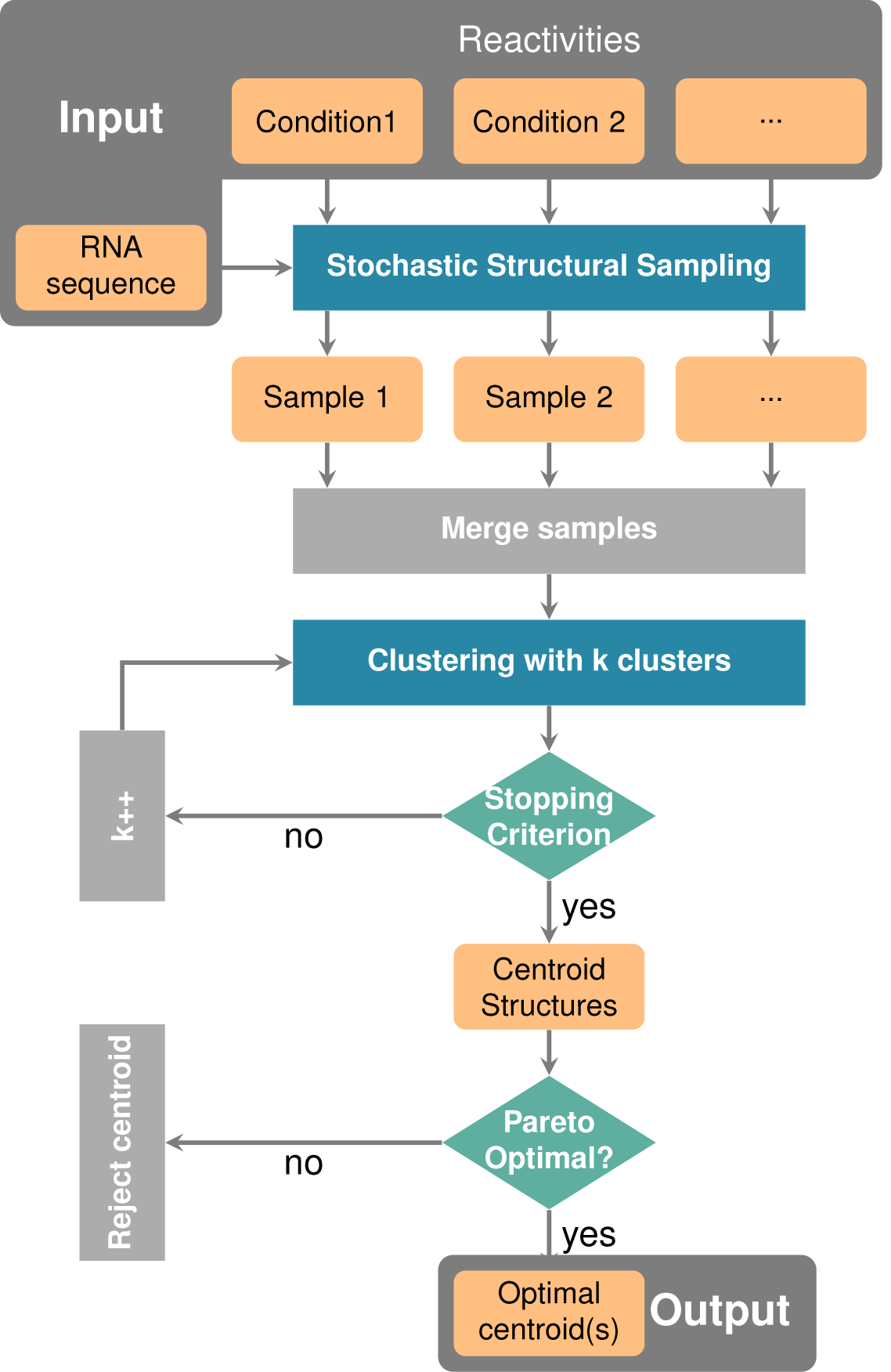
`IPANEMAP` workflow: `IPANEMAP` takes as input an RNA sequence with profiling data, denoted by reactivities, from various experimental conditions. `IPANEMAP` proceeds, first, with a stochastic sampling that results into samples of predicted secondary structure. The data-driven predicted structures are then gathered in one sample, serving as input for the clustering step. `IPANEMAP` proceeds, then, with an iterative clustering that ends once a stopping criterion is reached. This step allows to identify the adequate number of clusters *k* to be considered. The *k* resulting clusters are then represented by their centroid structures. Clusters figuring on the 2D-Pareto frontier are considered to be optimal and subsequently their corresponding centroids are reported as the predicted structure through `IPANEMAP`.

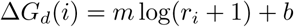

using *m* = 1.3 and *b* = −0.4. Those values were halved in comparison to those recommended by Deigan et al. (Deigan et al., 2009), following a grid search optimization on the Cordero dataset, and based on the rationale that lower absolute values for pseudo-energy bonuses increase the expected overlap between pseudo-Boltzmann ensembles. Those pseudo energy contributions effectively guide our predictions towards a subset of structures that are in good agreement with probing data. For each condition *d* in *𝒟*, we use the soft constraints framework (Lorenz et al., 2016) of the `RNAfold` software (Lorenz et al., 2011) to produce a random (multi)set 𝒮_*d*_ of *M* structures in the pseudo-Boltzmann ensemble.

#### Clustering across conditions

In order to infer recurrent conformations across sampled sets, `IPANEMAP` agglomerate structure (multi)sets while keeping track of their condition of origin, and *clusters* with respect to the *base-pair distance*, the number of base pairs differing between two structures. A clustering algorithm then partitions the (multi)set of sampled structures into *clusters*, (multi)sets of structures such that the accumulated sum of distances over clusters is minimized.

Among the many available options, we chose the Mini Batch *k*-Means algorithm (Sculley, 2010) (MBkM), implemented in the `scikit-learn` pacakage (Pedregosa et al., 2012), which requires less computational resources than the classic *k*-means algorithm, yet performed similarly in preliminary studies as an extensive collection of both agglomerative (affinity propagation) and hierarchical (Ward, Diana, McQuitty) clustering algorithms. A dissimilarity matrix, presenting the pairwise base-pair distance between structures, is precomputed and fed to the clustering algorithm.

Any cluster *C* output by the clustering is a multiset of structures, each labeled with its origin condition of *𝒟*. The *cluster probability of a structure feature f* (base pair or unpaired base) within a cluster *C* is then defined as

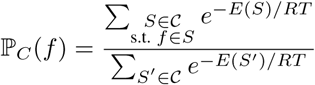

where *R* represents the Boltzmann constant, *E*(*S*) is the Turner free-energy, and *𝒞* is the (non-redundant) set of structures in *C*. From those probabilities we define the *centroid structure* of a cluster as its Maximum Expected Accuracy (MEA) structure, computed efficiently following Lu et al. (Lu et al., 2009).

Moreover, define the (pseudo-)*Boltzmann condition probability* of a structure *S*, generated for a probing condition *d* as part of a sampled set *𝒮*_*d*_, as

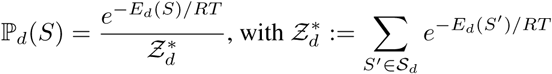

where *E*_*d*_(*S*) is the pseudo free-energy, including the Turner free-energy, assigned to the structure *S* within the probing condition *d*. The *stability* of a cluster *C* denoted its accumulated pseudo-Boltzmann probability across conditions, computed as

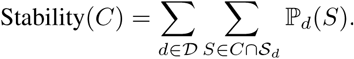

A cluster is deemed *significantly populated* if its stability exceeds a predefined threshold *E*. We set *E* = |*𝒟*| */*3 by default, such that at most three clusters are deemed significantly populated, and used as our primary candidates. Finally, we consider two clusters to be *highly similar* if their centroid structures differ by at most *δ* base pairs (*δ* = 1 by default), allowing the identification of clusters in the presence of minor variations.

The targeted number of clusters is a critical parameter of the MBkM algorithm. It should, at the same time, remain small enough to ensure reproducibility, while being sufficiently large to discriminate outliers and ensure consistency within each cluster. We determine an *optimal number of clusters k* using an iterative heuristic, gradually increasing the number of clusters until a significantly-populated cluster is split into two similar clusters, or a poorly populated (outlier) cluster is created. Namely, our iterative heuristic consists in running MBkM over an increasing number *k* of clusters, starting with *k* = 2, until the following *stopping criterion* is met: 1) Two significantly populated clusters have associated centroids which are highly similar; or 2) Centroid structures of significantly populated clusters from the previous iteration are highly similar to those of the current iteration.

#### Filtering the promising conformer(s)

Finally, we *select the most promising cluster(s)*, and return their centroid(s). While the final number of clusters may potentially be large, only a handful of clusters are expected to represent structures that are both stable, and supported by a large number of conditions. The remaining clusters are indeed probably artifacts of the clustering method, but nevertheless useful to filter out *noisy* structures.

We postulate that a perfect cluster should have large stability, as defined above, and be representative of several conditions. In the presence of a set of experimental conditions *𝒟*, we consider that a cluster *C* supports a given condition *d* when its probability within *d* exceeds a given threshold *τ*. The number of conditions supporting a cluster *C* is defined as

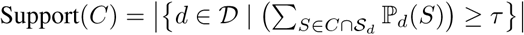

The value of *τ* is set to 1*/*(*k* + 1) with *k* the final number of clusters, ensuring at least one supporting cluster for each condition.

`IPANEMAP` evaluates the Stability and Support metrics for each cluster, and filters out any cluster that is dominated by some other with respect to both metrics. The remaining ones are *Pareto optimal*, a classic concept in multi-objective optimization (Mattson and Messac, 2005). `IPANEMAP` computes and returns the MEA centroid (Lu et al., 2009) of the Pareto-optimal clusters as its final prediction(s).

### Pairwise comparison of structural ensembles induced by reactivity profiles

We want to compare the structural ensembles induced by reactivity profiles, produced across diverse experimental conditions. To that purpose, we simply consider the base pair probability matrices, or dot-plots, resulting from supplementing the Turner energy model with pseudo energy terms. Dot plots can be computed efficiently in the presence of pseudo-energy terms using a variant of the McCaskill algorithm (McCaskill, 1990).

As a measure of the *ensemble distance* Dist induced by probing data, we consider the dot-plots associated with experimental conditions *d* and *d*′, and compute the squared Euclidean distance, such that

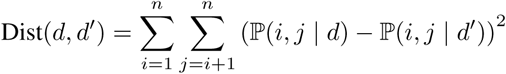

with *n* the length of the RNA sequence, ℙ (*i,j | d*) the Boltzmann probability of forming a base pair (*i, j*) in the pseudo-Boltzmann ensemble associated with condition *d*.

Individual dot-plots were computed using the `RNAfold` software in the Vienna Package 2.2.5, using the –p option in combination with the pseudo-energy terms introduced by Deigan et al. (Deigan et al., 2009).

### Datasets

To validate our computational method quantitatively, we consider several datasets, depending on the availability of probing data for one or several reagents, restricted to the wild type or produced for several point-wise mutants. Each dataset consists of sequences and individual reactivities to one or several probes, at each position in the RNA, completed with one or several functionally-relevant secondary structures.

#### Hajdin dataset

A dataset was gathered by Hajdin et al. (Hajdin et al., 2013) to validate the predictive capacities of probing data-driven predictions. It consists of 24 RNA sequences with known secondary structures for which a single chemical probing reactivity profile (1M7-SHAPE) was made available. This dataset includes sequences originating from a variety of organisms, and spans lengths ranging from 34 nts to 530 nts, with a focus on riboswitches and complex RNA architectures (full list in Supp. Table S3).

#### Cordero dataset

Probing data were downloaded from the RMDB (Cordero et al., 2012b) on July 2017. In the RMDB, reactivity scores are reported for all nucleotides, including those that are not expected to react with a given reagent. Thus, for the DMS (resp. CMCT) probing, we restricted reactivities to positions featuring nucleotides A and C (resp. G and U), setting the pseudo-energy term to 0 kcal.mol^-1^ for other positions. This allowed to decrease the noise generated by reactivities associated with non-targeted nucleotides, leading to more accurate predictions (data not shown).

#### Didymium structural model and probing data (DiLCrz dataset)

We considered the 188 nucleotides Lariat capping ribozyme from *Didymium iridis*, resolved 3.85 Å resolution using X-ray cristallography (PDB: 4P8Z) (Meyer et al., 2014). We annotated the secondary structure elements using the DSSR software from the 3DNA suite (Lu et al., 2015). Non-canonical base pairs were removed, and a non-pseudoknotted secondary structure was extracted as the maximum subset of non-crossing base pairs (Smit et al., 2008).

Probing data were experimentally generated, as described in the next section, using a comprehensive set of conditions covering some of the popular probing reagents and SHAPE technologies. We also considered the presence/absence of Mg^2+^, both to assess the capacity of `IPANEMAP` to recover tertiary interactions, and to assess the induced discrepancy on probing profiles and pseudo-Boltzmann ensembles.

#### Cheng dataset

Starting from the assumption that a functional structure should be preserved during evolution, we wanted to assess the agreement that might exist between probing data profiles for a set of RNA mutants. We considered DMS probing data, generated by (Cheng et al., 2017) through systematic point-wise mutations, for the Lariat-capping ribozyme (equivalent to 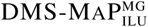 in the nomenclature below). We renormalized each reactivity profile following the method introduced by Deigan et al. (Deigan et al., 2009), restricted to the primer-free sequence: values greater than 1.5 times the interquartile range were discarded, and remaining values were divided by the mean of the top 10% reactivities. Overall, this constitutes a collection of 188 sequences, each having its associated reactivity profile.

### Experimental probing protocols

To systematically assess the potential of multiple sources of probing, we considered a difficult example, the *Didymium iridis* Lariat Capping ribozyme (DiLCrz). The native structure of DiLCrz, shown in Figure 4 is highly complex, and features two pseudoknots which cannot be explicitly modeled by most computational methods, making DiLCrz a challenging target for secondary structure prediction.

**Figure 2:**
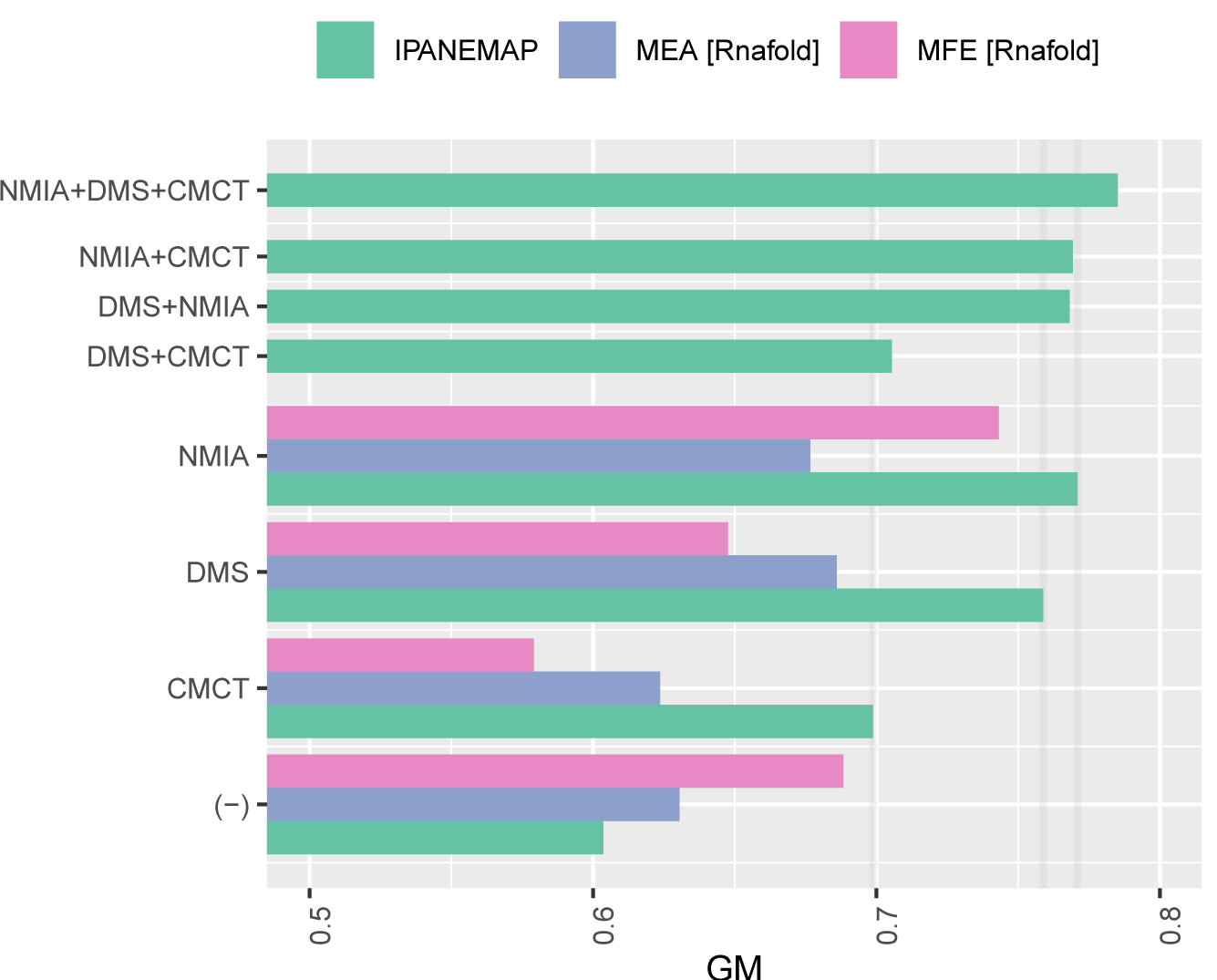
Predictive performances (GM) of `IPANEMAP` on the Cordero (Cordero et al., 2012b) dataset consisting of reactivities associated with the CMCT, NMIA and DMS probing for 6 reference RNAs. (Pseudo) MFE and MEA structures, predicted using `RNAfold` (Lorenz et al., 2011), are included as references in the case of a single condition, and in the absence of probing.

**Figure 3:**
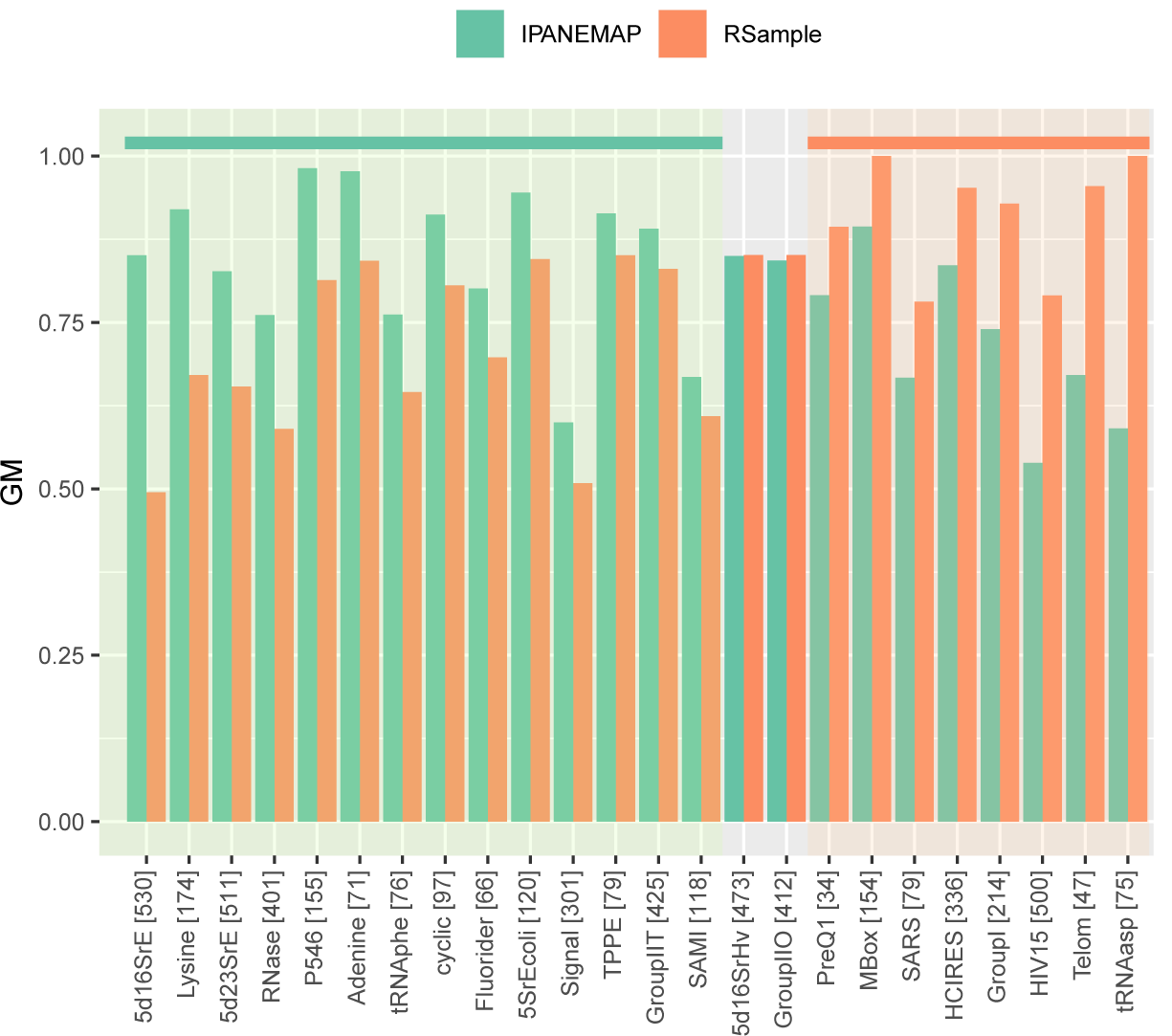
Comparison of predictive performances (GM) achieved by `IPANEMAP` and Rsample over the Hajdin et al. (Hajdin et al., 2013) data set, consisting of 24 RNAs which associated SHAPE reactivities. The length of each individual RNA is indicated within square brackets.

**Figure 4:**
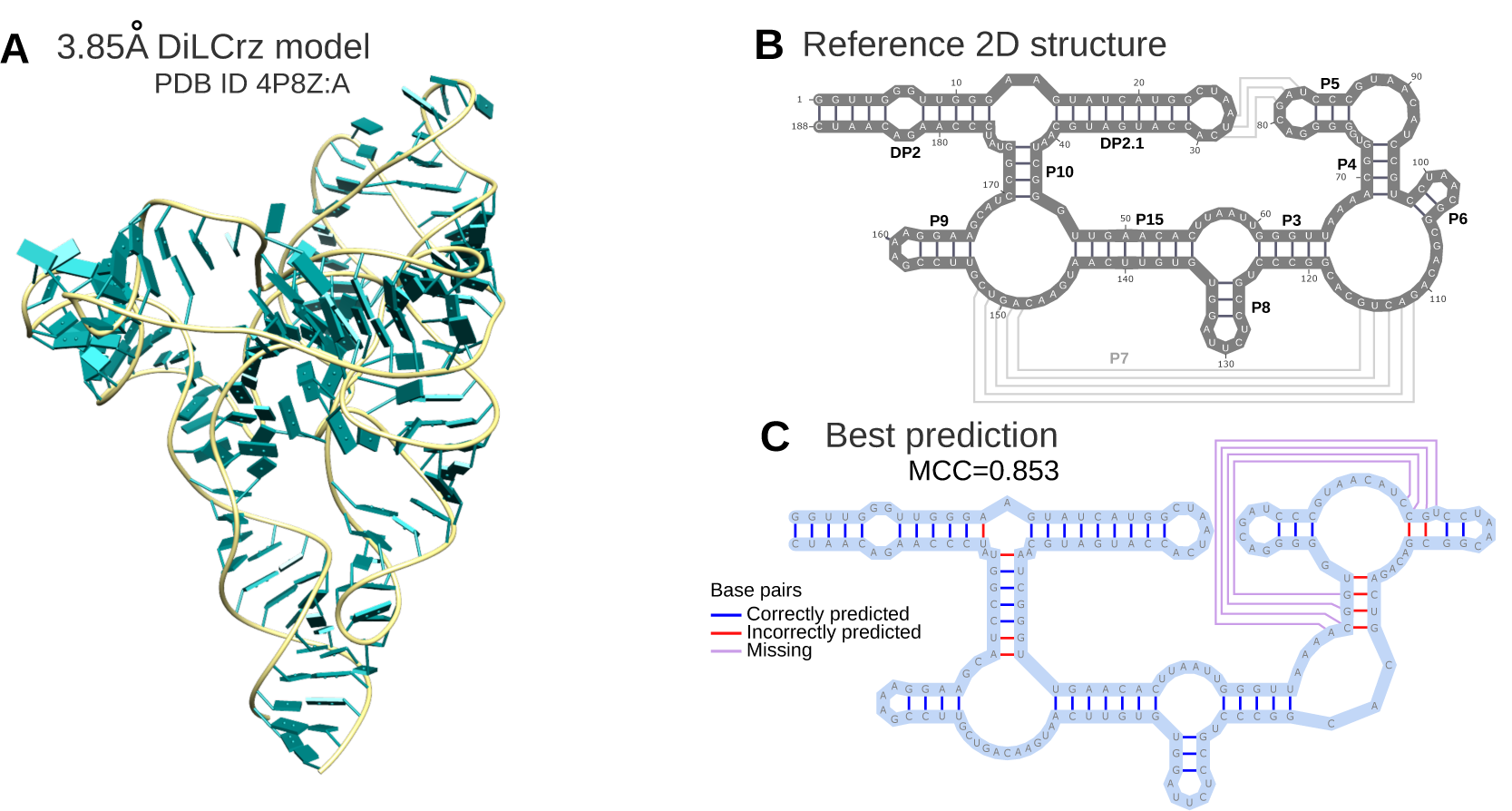
Model of the Lariat capping ribozyme from *Didymium iridis* (A – PDB 4P8Z:A by Meyer et al. (Meyer et al., 2014)), secondary structure, as annotated by DSSR (Lu et al., 2015) (B) and `IPANEMAP` first-ranking prediction (C.), obtained from the integration of 8 maximally-diverging probing conditions (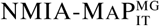, 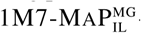-3D, NAI-CE^MG^, 1M7-CE, CMCT-CE^MG^,1M7-MAP_IL_ -3D, and 1M7-CE^MG^).

DiLCrz was probed with different SHAPE reagents: 1M7 (1-methyl-7 nitrosatoic anhydre), NMIA (N-Methyl Isatoic Anhydre), BzCN (Benzoyl cyanide) and NAI (2-methylnicotinic acid imidazolide) in presence and absence of Mg^2+^. DiLCrz was also probed with DMS (DiMethylSulfate) and CMCT, in presence of Mg^2+^, resulting in a total of 16 *probing conditions*, a term we use in the following to denote a combination of probing technology, reagent and presence/absence of Mg^2+^ (+ sequencing technology). For each probing condition, three experiments were performed in presence/absence of the reagent, and in a denatured context, following classic SHAPE protocols (Wilkinson et al., 2006; Smola et al., 2015).

As a preliminary experiment, we verified that DiLCrz generally adopts a single global architecture. To this end, we subjected DiLCrz to a standard denaturation/renaturation protocol (80°C for 2 min in H_2_O, addition of 40 mM of HEPES at 7.5 pH, 100 mM of KCl, 5 mM of MgCl_2_, followed by 10 min at room temperature, and 10 min at 37°C), and observed the production of a single band on a non-denaturing PAGE, strongly suggesting the adoption of a single conformation.

#### Stops-inducing probing protocol (SHAPE-CE)

6 pmol of RNA were resuspended in 18 *µ*l of water, denatured at 80°C for 2 minutes and cooled down at room temperature during 10 minutes in the probing buffer (40 mM HEPES at 7.5 pH, 100 mM KCl, in presence or absence of 5 mM MgCl_2_. After a 10 minutes incubation at 37°C, RNAs were treated with 2 mM of SHAPE reagent or DMSO (negative control) and incubated for 2 (BzCN), 5 (1M7), 30 (NMIA) or 60 (NAI) minutes at 37°C. Modified or unmodified RNAs were purified by ethanol precipitation and pellets were resuspended in 10 *µ*l of water. Modifications were revealed by reverse transcription using 5’ fluorescently labelled primers (D2 or D4 WellRED, Sigma Aldrich 5’ CTG-TGA-ACT-AAT-GCT-GTC-CTT-TAA 3’) and M-MLV RNAse (H-) reverse transcriptase (Promega) as previously described (Deforges et al., 2017), with only minor modification of the originally described SHAPE protocol (Wilkinson et al., 2006). cDNAs were separated by capillary electrophoresis (Beckman Coulter, Ceq8000). Data were analyzed using the `QuSHAPE` (Karabiber et al., 2013) software. RNA probing was performed in triplicate with distinct RNA preparations. The corresponding data sets are named after the probe followed by CE, and MG if probed in presence of MgCl_2_.

#### Mutations-inducing probing (SHAPE-MaP)

Probing was conducted as described in Smola et al. (Smola et al., 2015), except we used SuperScript III reverse transcriptase (ThermoFisher) at 50°C for 3 hours using the following specific primer:

5’ CTT-CAT-AGC-CTT-ATG-CAG-TTG-CTT-TTT-TTT-TTT TTT-TTT-TTT-GAT-TGT-CTT-GGG-ATA-CCG-GAT 3’

For 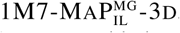, the denaturation step was repeated three times : After a 1 minute incubation at 95°C, 10 mM of 1M7 was added and the mix was incubated 1 minute at 95°C. This step was repeated 3 times for the samples labeled 3D. For NMIA-MAP_IT_ and 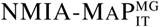, experiments were conducted as described above, except that the NGS library preparation was adapted to Ion Torrent sequencing. Sequences were mapped and analyzed with `ShapeMapper2` (Busan and Weeks, 2018) for 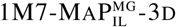 and 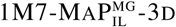. For the other conditions they are mapped with a script based on `ShapeMapper` (Smola et al., 2015), but adapted to Ion Torrent output files. The corresponding data sets are named after the probe followed by MAP. The presence of MG as a superscript indicates the presence of MgCl_2_, while the subscript IL (Illumina) or IT (Ion Torrent) specifies the sequencing technology.

#### DiMethylSulfate (DMS) and CMCT probing

DMS and CMCT probing were conducted essentially as described in (Weill et al., 2004; James and Sargueil, 2008). Succinctly, 6 pmol of RNA were resuspended in 18 *µ*l of water, denatured at 80°C for 2 minutes and cooled down at room temperature for 10 minutes in a probing buffer for DMS probing (40 mM HEPES pH 7.5, 100 mM KCl, 5 mM MgCl_2_) or in CMCT (50 mM of potassium borate pH 8, 100 mM KCl and 5 mM MgCl_2_). For DMS, RNA was then treated with 1 *µ*l of DMS (1:12 in ethanol) or 1 *µ*l of ethanol for 5 minutes at 37°C (mock reaction), the reaction was stopped by addition of 400 mM of Tris at 7.5 pH and immediately put on ice. For CMCT, RNA was treated with 10 *µ*l of CMCT (42 mg/mL) or 10 *µ*l H_2_O for 10 minutes at 37°C (mock reaction). Modified RNA were ethanol-precipitated and resuspended in 10 *µ*l of water. Modifications were mapped as described above for SHAPE-CE experiments.The corresponding data sets are named after the probe, followed by CE and MG.

### Benchmarking methodology

The Matthews Correlation Coefficient (MCC) is a classic metric for assessing the quality of a predicted structure *S*, identified by a set of base pairing positions, compared to an accepted reference (native) structure *S*^*\**^. It represents a compromise between the main metrics derived from the confusion matrix, and is defined as

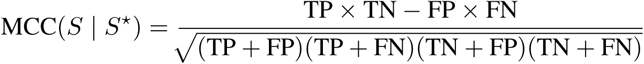

where TP, FP, TN, FN represent the correctly/erroneously predicted and correctly/erroneously omitted base pairs in *S* with respect to *S*^*\**^. MCC is a stringent metric, taking values between -100% and 100%, 0% being the expected MCC of a “coin tossing” random predictor.

For the sake of direct comparison with some competing methods (Spasic et al., 2017), we also report the Geometric Mean (GM) metric, defined as:

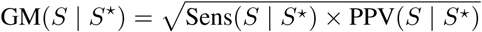

where Sens(*S* | *S*^*\**^) represents the proportion of base pairs in *S*^*\**^ that are in *S*, and PPV(*S* |*S*^*\**^) is the proportion of base pairs in *S* that are also in *S*^*\**^. This quality metric has very high correlation with the MCC, to which it is equivalent for large values of *n*, as shown by Gorodkin et al. (Gorodkin et al., 2001).

### Assessment of statistical significance

Following Xu et al. (Xu et al., 2012), we assessed the statistical significance of observed changes in predictive performances using a two-tailed Student *t*-test (unequal variance) with usual type I error rate *α* = 5%. When comparing performances within a given dataset, we used the paired version of the test.

Intuitively, the *t*-test estimates the probability that an observed change in performance can be attributed to chance alone. It considers the hypothesis *H*_0_ that the two distributions have equal mean and computes a *P-value, i*.*e*. the likelihood of the observed data under *H*_0_. If the P-value is lower than *α*, then *H*_0_ is deemed *refuted*, and the observed difference in performances is considered significant.

## Results

### Implementation

`IPANEMAP` was implemented in `Python 2.7+`, and mainly depends on the `RNAsubopt` software in the Vienna package (Lorenz et al., 2011), and `scikit-learn` (Pedregosa et al., 2012). `IPANEMAP` is free software distributed under the terms of the MIT license, and can be freely downloaded, along with a collection of helper utilities, from:

~~~
https://github.com/afafbioinfo/IPANEMAP
~~~

### Considering multiple probing conditions appears to improve the quality of predictions

Since `IPANEMAP` supports any number of probing profiles, we considered the Cordero et al. (Cordero et al., 2012b) dataset, further described in the previous section. It consists of 6 RNAs of known structures, for which reactivity profiles are available for the CMCT, NMIA and DMS reagents. We executed `IPANEMAP` with default parameters on each sequence and any subset of the three available conditions. The centroid secondary structure associated with the largest probability cluster was considered as the final prediction. Predictions were evaluated in term of the geometric mean (GM) metric, for the sake of comparison with previous studies (Spasic et al., 2017). As a control experiment, we also report `RNAfold` predictions in presence/absence of probing data, both in energy minimization (MFE) and maximum expected accuracy (MEA) modes. Figure 2 shows the averaged GM values for all combinations of tools and probing data (Detailed results in Supp. Tables S1 and S2).

In the absence of probing data, MFE predictions are generally dominant on this dataset, trailed by the MEA, and followed by `IPANEMAP` which, in this setting, devolves into the classic Ding-Lawrence algorithm (Ding and Lawrence, 2003). However, whenever probing reactivities are available, the single centroid returned by `IPANEMAP` always achieves higher GM values (Avg: 70%) than both MEA (Avg: 62%) and MFE (Avg: 58%), whose relative performances depend on the probing reagent. Interestingly, the quality of MFE and MEA predictions does not systematically benefit from additional probing data. Indeed, for half of the reagents and methods, the average GM obtained in the presence of a single reagent is lower than in the absence of probing data. Also, the impact of single probing data varies greatly across the three reagents, and the GE values of predictions respectively informed by CMCT, DMS and NMIA are ordered increasingly for all approaches, except for a minor reversal of DMS and NMIA in MEA mode.

The joint analysis of pairs of probing conditions appears to average the quality of predictions. Any combination of probing data yields a GM value that is always greater than the worst-performing condition in the pair, yet worse than the best-performing alone. Interestingly, the addition of the worst performing condition (CMCT), does not equally affect the performances of DMS (76% → 70.5%) and NMIA (77.1% → 76.9%), despite the latter conditions inducing similar GM values when considered alone. Indeed, CMCT+DMS yields GM values that are only remotely better than the worst-performing CMCT alone (70% → 70.5%), while CMCT+NMIA greatly outperforms CMCT alone (70% → 76.9%), almost matching the performance of the NMIA alone (77.1%). It is also worth mentioning that NMIA+CMCT, combining the best and worst conditions, achieves a better combined performance than DMS+NMIA, the two best mono-probing conditions.

Remarkably, the combination of the three conditions leads to the best overall predictions, averaging 78.5% GM. This seemingly improves by 8.5%, 2.7% and 1.4% over the average predictions achieved using CMCT, DMS and NMIA respectively. Table 1 illustrates the incremental refinement of `IPANEMAP` predictions for a glycine riboswitch upon increasing the number of considered probing conditions. However, these observations merely suggest an improvement, as statistical significance cannot be established on such a small dataset. Still, they suggests some level of complementarity between probing conditions, as already suggested by recent analyses (Brunel and Romby, 2000; Steen et al., 2012; Rice et al., 2014; Yu et al., 2018) and supported by further analyses in this paper.

**Table 1:**
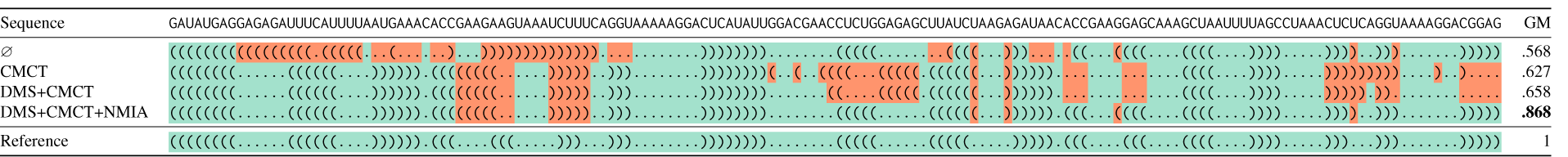
Dot-bracket representations of the native structure and `IPANEMAP` predictions for the glycine riboswitch sequence of the Cordero et al. (Cordero et al., 2012a) data set, supplemented with an increasing collection of probing conditions. Please note that, throughout the manuscript, by “conditions” we mean a combination of the nature of the probe, the probing technology, the ions present, the temperature, or any other “environmental” variation Positions with green and red backgrounds indicate correctly and incorrectly predicted base pairs.

### IPANEMAP performs comparably to state-of-the-art predictive methods when informed by a single profile

We assessed the predictive capacities of `IPANEMAP` in a classic setting where a single probing condition is available, and compared its performances against `RSample` software recently introduced by Spasic et al. (Spasic et al., 2017), `RSample` relies on a sampling/clustering method developed independently from the current work. It was shown to perform comparably as a comprehensive collection of state-of-the-art methods in probing-guided structure prediction, including `RME` (Wu et al., 2015), `RNAprob` (Deng et al., 2016), `RNAprobing` (Washietl et al., 2012), `RNAsc` (Zarringhalam et al., 2012) and the `fold` utility from the `RNAStructure` suite (Reuter and Mathews, 2010). We considered the Hajdin dataset (Hajdin et al., 2013), which consists of 24 RNAs of lengths ranging from 47 to 500 nts, all believed to fold into a unique documented conformation.

We used the default numbers of sampled structures for `RSample` and `IPANEMAP`, namely 10 000 to compute correction factors and 1 000 for the clustering phase. For all RNAs, a single structure was returned by both software. Note that the Pareto-optimality always implies a single returned cluster/structure for `IPANEMAP` in a mono probing setting. We computed and report in Figure 3 the individual Geometric Means (GM) of predicted structures. Sensitivity and PPV can be found in Supp. Table S4, while Supp. Table S5 reports the stability of predictions over 10 independent runs of `IPANEMAP`.

`IPANEMAP` appears to perform similarly, or even slightly favorably in comparison with `RSample`. It averages 80.1% GM, and reports more accurate structural models than its competitor for 14 RNAs in the Hajdin dataset (Hajdin et al., 2013), returns structures equally as good on 2 RNAs, and is dominated by `RSample` for 8 RNAs. However, its modest average improvement of approximately 1.5% over the dataset is not statistically significant, according to a two-tailed paired T-test. Those results support the notion that `IPANEMAP` represents a competitive alternative to state-of-the-art methods for single reactivity profiles.

### Comparing and exploiting multiple probing conditions: A case study on the Lariat capping ribozyme

To assess the potential of our method, we considered the Lariat capping ribozyme from *Didymium Iridis* (DiLCrz) whose 3D structure has been modeled at 3.85 Å from X-ray diffraction data (Meyer et al., 2014). This RNA contains a large diversity of interactions, including helical domains, tertiary pairings, a kissing loop and a pseudoknot (Figure 4). Its automated modeling is therefore a challenging case for computational structure prediction methods. For instance, running `RNAfold` with default parameters only recovers four (DP2, DP2.1, P10 and P9) of the ten helices. Those are universally recovered by energy-based prediction tools, so differences in performances will mainly be observed between nucleotides 35 and 163.

As described in the Material and Methods section, we probed DiLCrz in 16 different conditions using different SHAPE and classical reagents, in presence/absence of Mg^2+^ ions, using either the standard premature stop or the newly developed mutation induction technologies to map the modification sites. We observed an overall good agreement of reactivities with the 3D model proposed by Meyer et al. (Meyer et al., 2014) (see Supp. Figures S1, S2 and S3). Reactivities were then used as input of `IPANEMAP`, either individually or combined, for secondary structure prediction.

### Comparison and clustering of conditions

Faced with diverse probing conditions, we first assessed the compatibility of the conclusions drawn from different probing data, including the probing-free run of `IPANEMAP` as a control. We used the methodology described in the Material and Methods and, for each condition, computed the base pair probability distribution (aka *dot-plot*) in the pseudo-Boltzmann ensemble induced by the reactivity profile. We then computed the Ensemble Distance, the squared Euclidean distance between dot plots, for each pair of conditions. Figure 5 summarizes the pairwise distance, projected onto a 2D surface by a principal component analysis. A systematic cross-analysis of raw reactivities revealed a positive, if modest, correlation between conditions, but did not support a clear partition of profiles based on probing technology or presence/absence of Mg^2+^ (see Supp. Figure S4).

**Figure 5:**
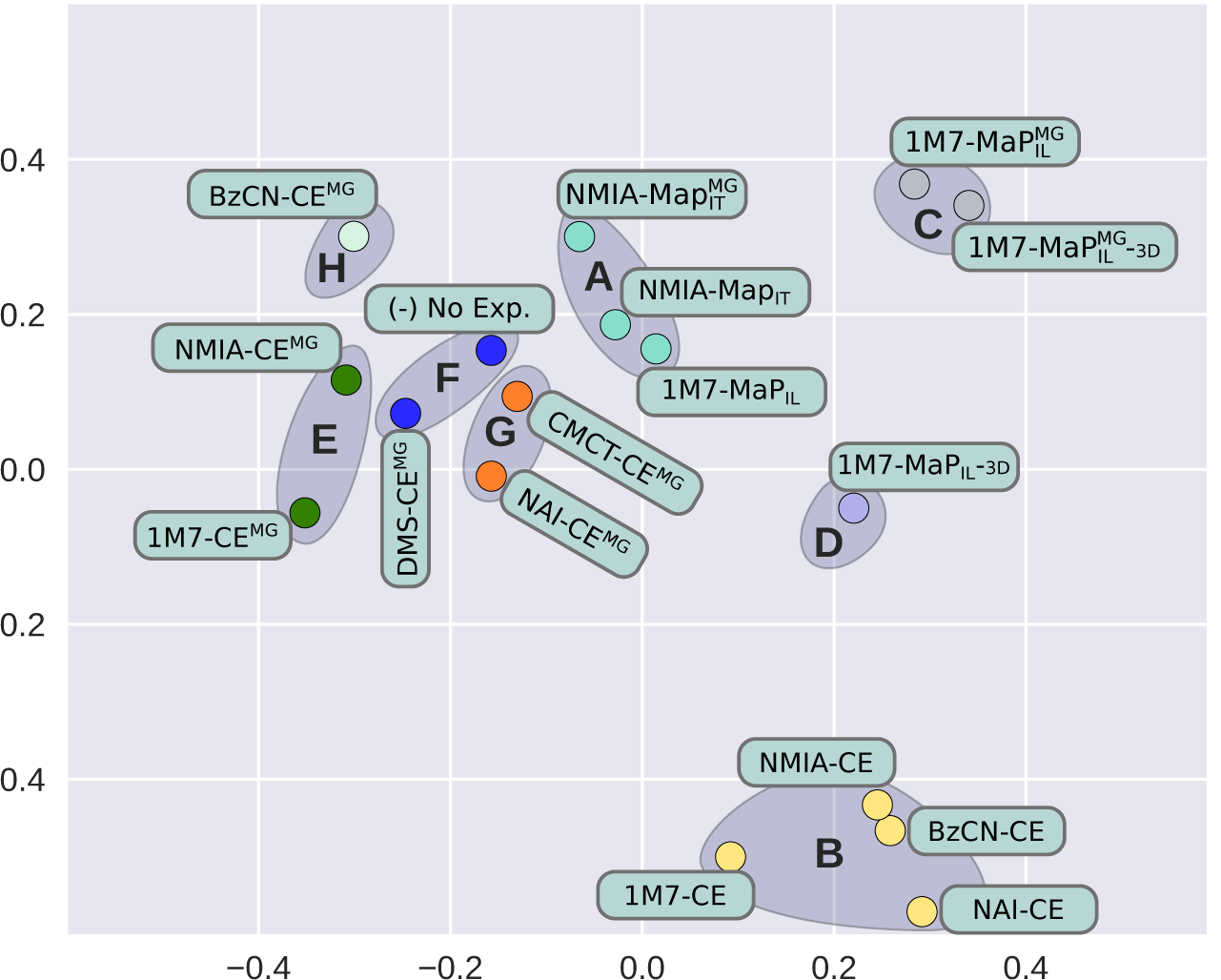
2D spatial representation of conditions, computed by Principal Component Analysis (PCA) to optimally reflect pairwise ensemble distances between conditions. Colors and grayed areas indicate clusters of conditions.

A visual inspection of Figure 5 suggests the presence of 8 clusters. In order to objectively build groups of compatible conditions, we performed a k-mean clustering using `scikit-learn`, setting k to 8, and obtained the clusters highlighted in Figures 6 and 5. We obtain the following clusters:

**Figure 6:**
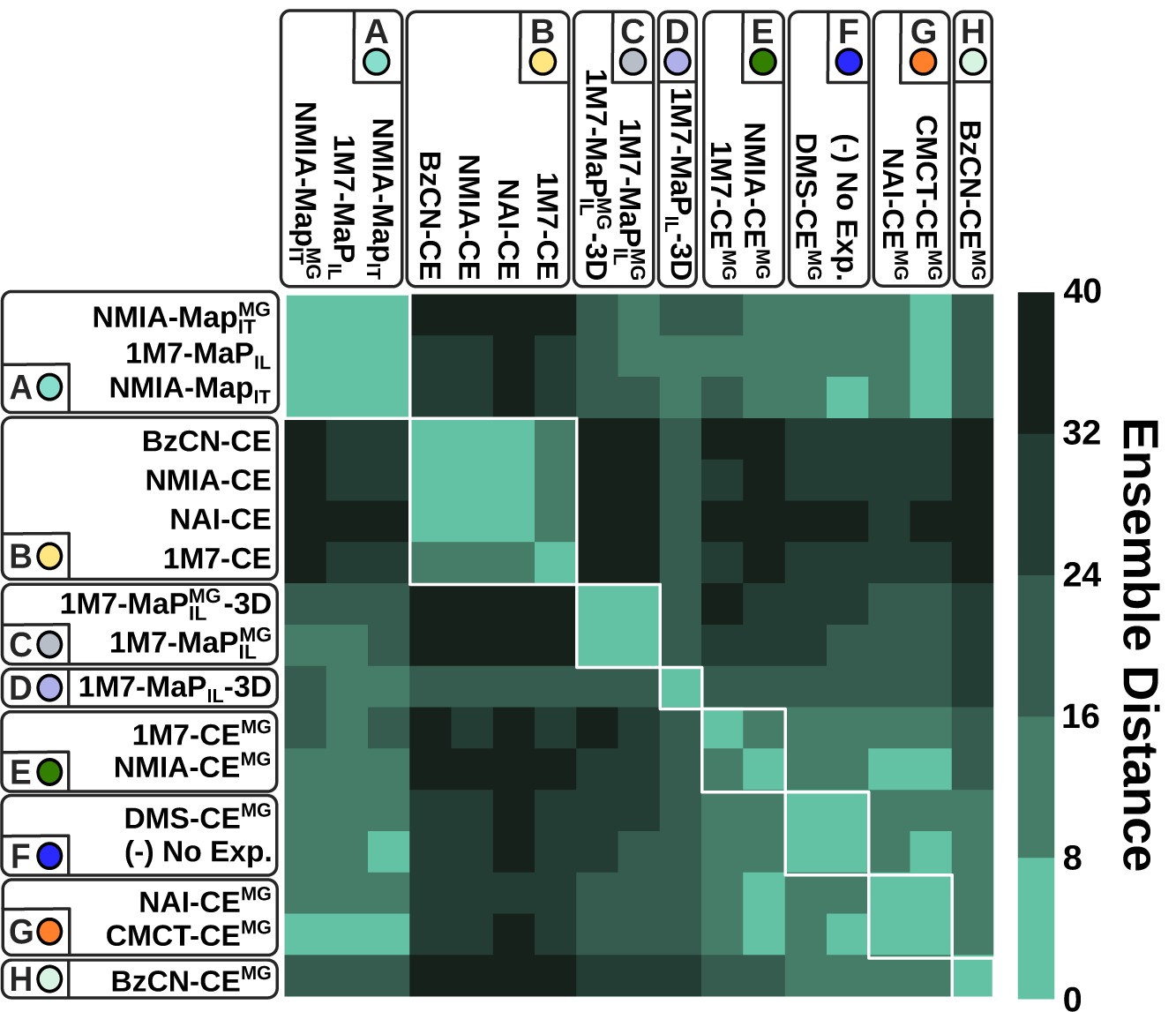
Discretized ensemble distance induced by conditions. Conditions ordered by spectral biclustering. Conditions regrouped within clusters by k-mean clustering appear as blocks on the x and y axis.

- A 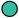: 1M7-MAP_IL_, NMIA-MAP_IT_, and 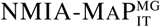;
- B 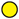: 1M7-CE, NMIA-CE, BZCN-CE, and NAI-CE;
- C 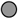: 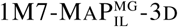 -3D and 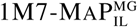;
- D 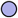: 1M7-MAP_IL_ -3D;
- E 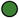: NMIA-CE^MG^ and 1M7-CE^MG^;
- F 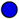: DMS-CE^MG^ and probing-free;
- G 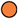: CMCT-CE^MG^ and NAI-CE^MG^;
- H 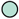: BZCN-CE^MG^.

The predicted clusters are generally consistent with the ordering resulting from a spectral biclustering, implemented in `scikit-learn` and executed with default parameters, as illustrated by Figure 6.

The average ensemble distance and PCA visualization support a status of outliers for conditions in the cluster B 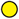 (1M7-CE, NMIA-CE, BZCN-CE, and NAI-CE). We hypothesize that such conditions may be either representative of alternative conformations, or be altogether erroneous. Consequently, we will single out this cluster within our detailed analysis of the multi-probing performances.

#### Assessment of predictions informed by individual probing conditions (mono probing)

For each probing condition, we executed `IPANEMAP` on a single reactivity profile, using a sample size of 1 000 structures. For the sake of reproducibility, we executed `IPANEMAP` 10 times for each condition, and report in Table 2 the average MCC values. We also report for reference the MCCs obtained by running `RNAfold` with default parameters in energy minimization (MFE) and Maximum Expected Accuracy (MEA) modes.

**Table 2:**
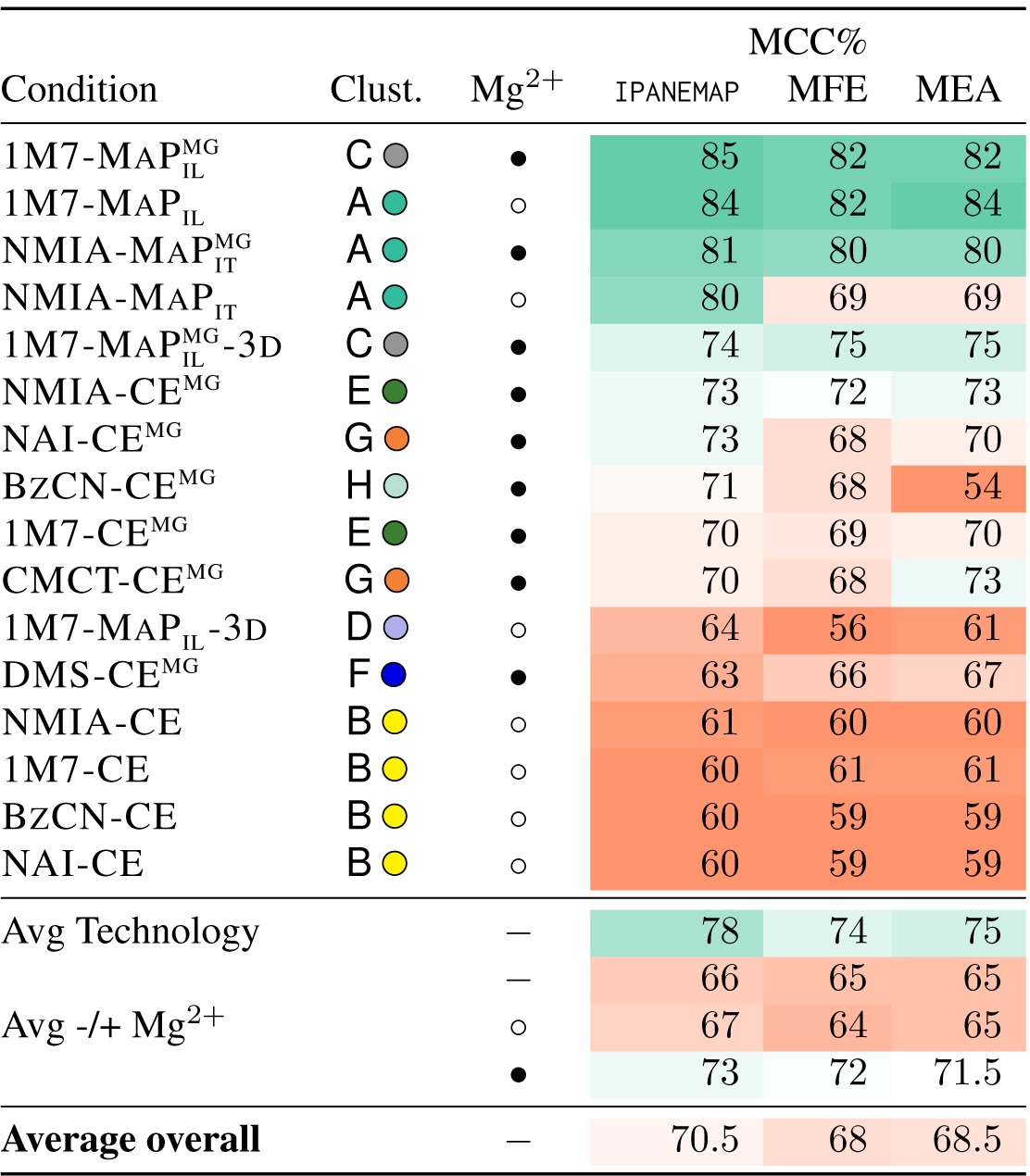
Predictive performances (MCC) of `IPANEMAP`, averaged over 10 runs, for the DiLCrz dataset, consisting of 16 reactivity profiles. (Pseudo) MFE and MEA structures, predicted using `RNAfold` (Lorenz et al., 2011), are included for reference. The structure was probed in presence (•) or absence (○) of Mg^2+^.

The predictive performances of `IPANEMAP` averages a 70.5% MCC, compared to 68.5% and 68% average MCC for MEA and MFE predictions respectively. This suggests modest, barely significant, improvements compared to classic prediction paradigms (P-values of 2% and 14% respectively). The stability of predictions across conditions seem comparable for all `IPANEMAP`, MEA and MFE-driven predictions (std dev. of 8%). Excluding the outliers of cluster B 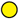 improved the average MCC to 74%.

Across the sixteen conditions, we observed a large discrepancy in the capacity of the experimental setup to inform predictions, with MCCs ranging from above 80% (NMIA-MAP_IT_, 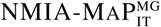, 1M7-MAP_IL_ and 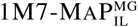) to around 60% (NAI-CE, BZCN-CE, 1M7-CE and NMIA-CE). Interestingly the MCC value cannot be trivially anticipated from the compatibility of the reactivity scores with the predicted structure, or even with the native structure (see Supp. Table S1). These predictive capacities clearly outperformed those achieved in the absence of probing data (51% MCC), and can be interpreted as indicative of predictions of good overall quality. The lowest MCC value of 60% is, in particular, equally consistent with Sensitivity/PPV values of 60%/60%, 80%/46% or 45%/82%.

Unsurprisingly, the presence/absence of Magnesium ions during the probing was observed to impact the predictions, with an observed drop from an average 73% MCC in the presence of Mg^2+^ to 67% MCC in the absence of Mg^2+^. However, while the poorly performing conditions in Cluster B 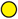 were probed in the absence of Mg^2+^, some of the most informative profiles (1M7-MAP_IL_ and NMIA-MAP_IT_) were obtained in absence of Mg^2+^. Even more drastic changes of performances were observed when comparing the MCCs of SHAPE-MaP experiments (avg 78%) with the historical SHAPE CE technology (avg 66%).

### Bi-probing mitigates the influence of outliers, but does not significantly improve prediction quality

Next, we turned to a systematic exploration of the predictive capacities of bi-probing analyses, based on pairs of probing profiles, and attempted to quantify their impact on `IPANEMAP` predictions. For each pair and triplets of conditions, we executed `IPANEMAP` using 1 000 structures per condition, and considered the first returned structure. We then computed the associated MCC, and compared it with the MCC of the worst performing condition (Min), best-performing condition (Max) and average MCC over the single conditions experiments. A summary of the results over pairs is reported in Figure 7 (details in Supp. Table S6).

**Figure 7:**
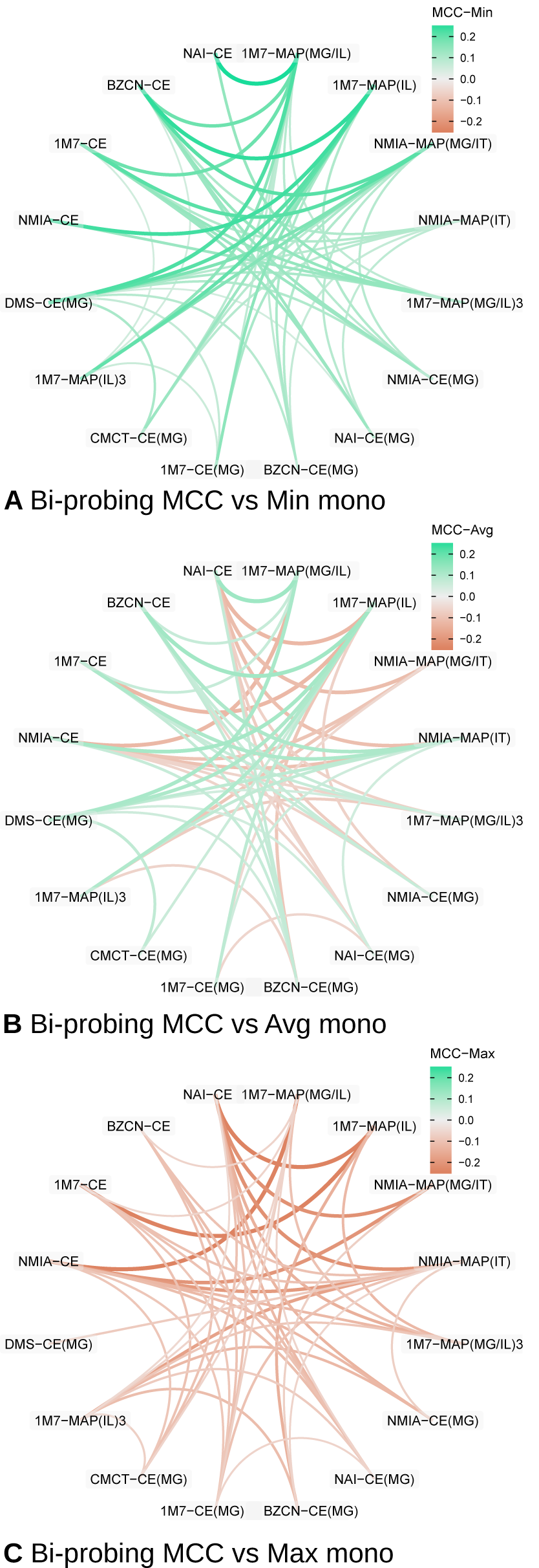
Shift in `IPANEMAP` predictive capacity (MCC) observed upon considering two conditions, compared to the worst (**A**), average (**B**) and best (**C**) predictions achieved for mono-probing conditions. Green and red edges indicate enhanced and degraded predictions respectively compared to the reference (Min, Max or Average of conditions). Thickness indicates absolute shift value (only values above 5% reported for readability).

A superficial inspection of the resulting MCCs appears to confirm the conclusions reached on the Cordero et al. (Cordero et al., 2012a) dataset. Indeed, considering two conditions led to predictions whose quality fall between the worst and the best one, while being generally close to the average. More precisely, the average MCC value of pair-informed predictions remained at 70.7%, improving only by 0.2% over the average MCC of mono-probing predictions. Pair-informed predictions were marginally more reliable, with the median MCC increasing to 72.5% MCC from 70.5% for mono-probing predictions. However, outliers strongly impacted the overall picture, and ignoring the conditions in cluster B 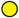 increased the average MCC to 75.1%, while unsurprisingly decreasing the standard deviation (8.6% → 6.5%).

Compared to the minimum MCC of the pair, the MCC of bi-probing predictions was improved by 5.3% on average over the MCC of the worst-performing conditions. Pair MCCs exceeded their associated min. MCC by at least 5% in 47 pairs out of the 120 possible pairs, while never being dominated by more than 5% MCC.

Bi-probing seemed to perform similarly as the average of the two mono-probing conditions, with an average MCC improvement of 0.2%. However, for 56 out of the 120 possible pairs, the bi-probing conditions exhibited an improvement of at least 1% over the average, while a decay of at least 1% was observed for 46 out of the remaining 64 pairs. Such a decay can partly be attributed to the disruptive effect of the outliers from cluster B 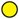, involved in 35 of the 46 observed loss of predictive performance. Moreover, removing this cluster induced an average MCC improvement of 1.1%.

Predictions informed by two conditions remained, however, generally dominated by the best performing condition of the pair. On average, the MCC of the best condition is 4.9% higher than the one achieved by bi-probing analysis. Outliers in cluster B 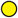 are partly responsible for this situation, and their removal reduces the average MCC decay to 3.3%. Moreover, for 9 pairs of conditions, the bi-probing analysis produced MCCs that are at least 1% better than the best of its two conditions, but these encouraging examples were generally dominated by 66 examples where the pair performs more than 1% worse than the best mono-probing condition.

#### Considering three conditions yields improvement over the average of mono conditions within the same triplet

Increasing the number of conditions to three boosted the average MCC to 72.3% and even reached 76.5% over triplets that do not contain conditions from cluster B 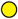. The MCC gain over the worst condition in the triplet reached 9.1% on average. Predictions were also more consistently good, with a median MCC of %73.4. Disregarding conditions in cluster B 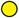 did stabilize the quality of prediction (stdev 8.4% → 5.5%), while increasing the average MCC to 77.4%.

Compared to bi-probing, MCC values achieved by integrating triplets of conditions indicated a clear gain over the average performances of individual conditions, supporting a notion of complementary between experiments. Indeed tri-probing experiments induced an average +1.7% MCC improvement (median +2.6) over the average of mono-probing prediction in the triplet (P-value= 1.4 *×* 10^−9^). The gain of quality was widespread, and observed for 65% of the triplets. Finally, 39 triplets were found to induce MCC values greater than 83%.

The enhanced performances of tri-probing over bi-probing was deemed statistically significant (P-value= 2 *×* 10^−2^), and consistent with the *voting* principle underlying the clustering used in `IPANEMAP`. Indeed, in a bi-probing setting, a single outlier (*e*.*g*. NAI-CE) may entirely determine the final structure due to its Boltzmann ensemble being very tightly concentrated around a, presumably erroneous, centroid structure. In the presence of three or more conditions, however, clusters resulting from an outlier condition are typically expected to be dominated in the Pareto front, by compatible structures originating from alternative conditions. It follows that the influence of outliers over the final prediction is mitigated.

#### Similar conditions do not contribute to the quality of predictions

Our distance assessment and clustering on conditions revealed groups of conditions that were highly compatible in their conclusions, while others seemed to include highly diverging structural information. While some outlier conditions, such as those of cluster B 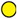, clearly stood out as erroneous, the results of our systematic analysis of pairs and triplets supported a notion of *complementarity between conditions*. Under this assumption, downstream analyses would be more likely to benefit from the presence of highly diverging probing experiments, than the accumulation of similar (arguably redundant) profiles.

To test this hypothesis, we executed `IPANEMAP` on every subset of conditions within clusters, and report in Table 3 the quality (MCC) of predictions for any subset of similar conditions (see Supp. Figure S5 for associated models). Note that such an analysis does not use knowledge of the native structure, since our clustering only relies on properties of the pseudo-Boltzmann ensemble.

**Table 3:** Predictions do not benefits from the joint consideration of similar conditions. For each cluster, all subsets of conditions in the cluster are considered, and the MCC of the resulting prediction (‘Joint’) is compared to the average MCC of predictions performed with individual conditions independently (‘Mono’), revealing little improvement.

|  | Conditions | MCC% |  |
| --- | --- | --- | --- |
|  |  | Joint | Mono |
| Cluster A ● | NMIA-MAP<sub>IT</sub><sup>MG</sup>, 1M7-MAP<sub>IL</sub>, NMIA-MAP<sub>IT</sub> | 80 | 82 |
|  | NMIA-MAP<sub>IT</sub><sup>MG</sup>, 1M7-MAP<sub>IL</sub> | 82 | 82 |
|  | 1M7-MAP<sub>IL</sub>, NMIA-MAP<sub>IT</sub> | 82 | 82 |
|  | NMIA-MAP<sub>IT</sub><sup>MG</sup>, NMIA-MAP<sub>IT</sub> | 80 | 80 |
| Cluster B ● | NAI-CE, BzCN-CE, 1M7-CE, NMIA-CE | 60 | 60 |
|  | NAI-CE, BzCN-CE, 1M7-CE | 60 | 60 |
|  | NAI-CE, BzCN-CE, NMIA-CE | 61 | 60 |
|  | NAI-CE, 1M7-CE, NMIA-CE | 61 | 60 |
|  | BzCN-CE, 1M7-CE, NMIA-CE | 61 | 60 |
|  | NAI-CE, BzCN-CE | 60 | 60 |
|  | NAI-CE, 1M7-CE | 59 | 60 |
|  | NAI-CE, NMIA-CE | 60 | 60 |
|  | BzCN-CE, 1M7-CE | 61 | 60 |
|  | BzCN-CE, NMIA-CE | 60 | 60 |
|  | 1M7-CE, NMIA-CE | 62 | 61 |
| Cluster C ● | 1M7-MAP<sub>IL</sub><sup>MG</sup>, 1M7-MAP<sub>IL</sub><sup>MG</sup>-3D | 82 | 80 |
| Cluster E ● | 1M7-CE<sup>MG</sup>, NMIA-CE<sup>MG</sup> | 73 | 72 |
| Cluster G ● | NAI-CE<sup>MG</sup>, CMCT-CE<sup>MG</sup> | 73 | 72 |

As expected, compared to the average MCC over associated mono-probing analyses, a joint consideration of similar conditions only induced limited progress over single probing analyses (between -1% and +1%), leading to an modest average improvement (+0.2% average MCC). This is consistent with the notion that supplementing a probing-based modeling with compatible conditions provides very little additional information, leading to a limited contribution of similar conditions.

#### Considering diverse conditions improves and stabilizes the quality of predictions

Having established the redundant nature of similar conditions, and their limited contribution to downstream modeling, we turn to an analysis of conditions across clusters. We executed `IPANEMAP` to assess the quality of predictions informed by conditions chosen across the 8 clusters. We considered any possible combination, except for the outlier cluster B 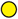, for which we only considered NAI-CE as a representative. The results, summarized in Table 4, demonstrate a clear advantage of including diverse conditions, with an average MCC value of 77% in the absence of condition from cluster B 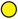. Those performances compare favorably against the 72% MCC achieved on average by individual conditions (P-value= 3 *×* 10^−5^), and reveal a remarkable stability (5% stdev), and a degradation in only two out the 24 tested cases (∼ 1% loss of MCC in both cases). A similar trend can be observed in the disruptive presence of NAI-CE, inducing an 72% average MCC, albeit with an increased standard deviation of 9%.

**Table 4:** Predictive performances of conditions across clusters. A single condition is chosen from each cluster except for B 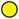. Clusters consisting of a single condition (D 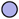, F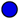 and H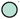) contribute additional conditions := {1M7-MAP_IL_ -3D, BZCN-CE^MG^, DMS-CE^MG^}. The MCC of the predicted structure is reported (‘Joint’ columns) and compared to the average of mono conditions (‘Mono’ columns), in presence/absence of NAI-CE.

| Conditions in individual clusters (+$\mathcal{M}$) | | | | MCC% | | | |
| --- | --- | --- | --- | --- | --- | --- | --- |
|  |  |  |  | -NAI |  | +NAI |  |
| A $\bullet$ | C $\bullet$ | E $\bullet$ | G $\bullet$ | Joint | Mono | Joint | Mono |
| NMIA-MAP<sub>IT</sub><sup>MG</sup> | 1M7-MAP<sub>IL</sub><sup>MG</sup> | 1M7-CE<sup>MG</sup> | NAI-CE<sup>MG</sup> | 84 | 72 | 83 | 71 |
| NMIA-MAP<sub>IT</sub><sup>MG</sup> | 1M7-MAP<sub>IL</sub><sup>MG</sup> | 1M7-CE<sup>MG</sup> | CMCT-CE<sup>MG</sup> | 75 | 72 | 82 | 71 |
| NMIA-MAP<sub>IT</sub><sup>MG</sup> | 1M7-MAP<sub>IL</sub><sup>MG</sup> | NMIA-CE<sup>MG</sup> | NAI-CE<sup>MG</sup> | 79 | 73 | 82 | 71 |
| NMIA-MAP<sub>IT</sub><sup>MG</sup> | 1M7-MAP<sub>IL</sub><sup>MG</sup> | NMIA-CE<sup>MG</sup> | CMCT-CE<sup>MG</sup> | 82 | 72 | 60 | 71 |
| NMIA-MAP<sub>IT</sub><sup>MG</sup> | 1M7-MAP<sub>IL</sub><sup>MG</sup>-3D | 1M7-CE<sup>MG</sup> | NAI-CE<sup>MG</sup> | 70 | 71 | 60 | 70 |
| NMIA-MAP<sub>IT</sub><sup>MG</sup> | 1M7-MAP<sub>IL</sub><sup>MG</sup>-3D | 1M7-CE<sup>MG</sup> | CMCT-CE<sup>MG</sup> | 70 | 70 | 72 | 69 |
| NMIA-MAP<sub>IT</sub><sup>MG</sup> | 1M7-MAP<sub>IL</sub><sup>MG</sup>-3D | NMIA-CE<sup>MG</sup> | NAI-CE<sup>MG</sup> | 72 | 71 | 74 | 70 |
| NMIA-MAP<sub>IT</sub><sup>MG</sup> | 1M7-MAP<sub>IL</sub><sup>MG</sup>-3D | NMIA-CE<sup>MG</sup> | CMCT-CE<sup>MG</sup> | 75 | 71 | 60 | 70 |
| 1M7-MAP<sub>IL</sub> | 1M7-MAP<sub>IL</sub><sup>MG</sup> | 1M7-CE<sup>MG</sup> | NAI-CE<sup>MG</sup> | 82 | 73 | 75 | 71 |
| 1M7-MAP<sub>IL</sub> | 1M7-MAP<sub>IL</sub><sup>MG</sup> | 1M7-CE<sup>MG</sup> | CMCT-CE<sup>MG</sup> | 84 | 72 | 60 | 71 |
| 1M7-MAP<sub>IL</sub> | 1M7-MAP<sub>IL</sub><sup>MG</sup> | NMIA-CE<sup>MG</sup> | NAI-CE<sup>MG</sup> | 73 | 73 | 60 | 72 |
| 1M7-MAP<sub>IL</sub> | 1M7-MAP<sub>IL</sub><sup>MG</sup> | NMIA-CE<sup>MG</sup> | CMCT-CE<sup>MG</sup> | 82 | 73 | 82 | 71 |
| 1M7-MAP<sub>IL</sub> | 1M7-MAP<sub>IL</sub><sup>MG</sup>-3D | 1M7-CE<sup>MG</sup> | NAI-CE<sup>MG</sup> | 84 | 71 | 60 | 70 |
| 1M7-MAP<sub>IL</sub> | 1M7-MAP<sub>IL</sub><sup>MG</sup>-3D | 1M7-CE<sup>MG</sup> | CMCT-CE<sup>MG</sup> | 70 | 71 | 84 | 70 |
| 1M7-MAP<sub>IL</sub> | 1M7-MAP<sub>IL</sub><sup>MG</sup>-3D | NMIA-CE<sup>MG</sup> | NAI-CE<sup>MG</sup> | 74 | 72 | 60 | 70 |
| 1M7-MAP<sub>IL</sub> | 1M7-MAP<sub>IL</sub><sup>MG</sup>-3D | NMIA-CE<sup>MG</sup> | CMCT-CE<sup>MG</sup> | 74 | 71 | 75 | 70 |
| NMIA-MAP<sub>IT</sub> | 1M7-MAP<sub>IL</sub><sup>MG</sup> | 1M7-CE<sup>MG</sup> | NAI-CE<sup>MG</sup> | 82 | 72 | 75 | 71 |
| NMIA-MAP<sub>IT</sub> | 1M7-MAP<sub>IL</sub><sup>MG</sup> | 1M7-CE<sup>MG</sup> | CMCT-CE<sup>MG</sup> | 83 | 72 | 82 | 70 |
| NMIA-MAP<sub>IT</sub> | 1M7-MAP<sub>IL</sub><sup>MG</sup> | NMIA-CE<sup>MG</sup> | NAI-CE<sup>MG</sup> | 73 | 73 | 59 | 71 |
| NMIA-MAP<sub>IT</sub> | 1M7-MAP<sub>IL</sub><sup>MG</sup> | NMIA-CE<sup>MG</sup> | CMCT-CE<sup>MG</sup> | 82 | 72 | 82 | 71 |
| NMIA-MAP<sub>IT</sub> | 1M7-MAP<sub>IL</sub><sup>MG</sup>-3D | 1M7-CE<sup>MG</sup> | NAI-CE<sup>MG</sup> | 73 | 71 | 82 | 69 |
| NMIA-MAP<sub>IT</sub> | 1M7-MAP<sub>IL</sub><sup>MG</sup>-3D | 1M7-CE<sup>MG</sup> | CMCT-CE<sup>MG</sup> | 70 | 70 | 73 | 69 |
| NMIA-MAP<sub>IT</sub> | 1M7-MAP<sub>IL</sub><sup>MG</sup>-3D | NMIA-CE<sup>MG</sup> | NAI-CE<sup>MG</sup> | 74 | 71 | 68 | 70 |
| NMIA-MAP<sub>IT</sub> | 1M7-MAP<sub>IL</sub><sup>MG</sup>-3D | NMIA-CE<sup>MG</sup> | CMCT-CE<sup>MG</sup> | 74 | 71 | 73 | 69 |
| Average |  |  |  | 77 | 72 | 72 | 70 |

Finally, considering the 16 conditions together induced a 80% MCC value, increasing to 83% in the absence of cluster B 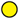. Overall, these results support the notion that the strategy used by `IPANEMAP` is able to exploit the complementary information provided by multiple conditions, contributing to better predictions.

### Supplementing wild-type profiles with random single mutants increases prediction quality

In this final analysis, we tested the capacity of `IPANEMAP` to extract and exploit complementary information produced by the systematic probing of single-point mutants using the Mutate-and-Map (MaM) protocol (Cordero and Das, 2015). The joint analysis of several mutants may appear error-prone, as single point mutants are expected to adopt different structures than the wild-type sequence (WT), an expectation that is at the core of downstream analyses (Cordero and Das, 2015). However, such changes are typically local, and the robustness of `IPANEMAP` to outlier conditions nourishes the hope that the benefits of including independently-produced conditions will outweigh the cost of noise introduced by local changes.

We generated 100 uniformly-distributed subsets consisting of 1, 2, 3 and 10 mutants extracted from the Cheng dataset (Cheng et al., 2017), which we each supplemented with the WT sequence. For each set of probing profiles 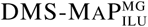, we executed `IPANEMAP` with default parameters, using a sample size of 1 000 structures. We also analyzed the WT sequence alone, reproducing the analysis 100 times to compare the variability across sets of mutants to the one induced by stochastic aspects of our method. Finally, we performed a restricted analysis focusing on the WT, supplemented with 2 probing profiles selected from the 20 most similar conditions to the WT, as assessed by the ensemble distance metric. By comparing our results with those obtained for unrestricted pairs of mutants, we assess the influence of structure variability on the quality of our predictions. We report in Figure 8 the distribution of MCC values associated with the first prediction returned by `IPANEMAP` (see Supp. Figure S6 for details).

**Figure 8:**
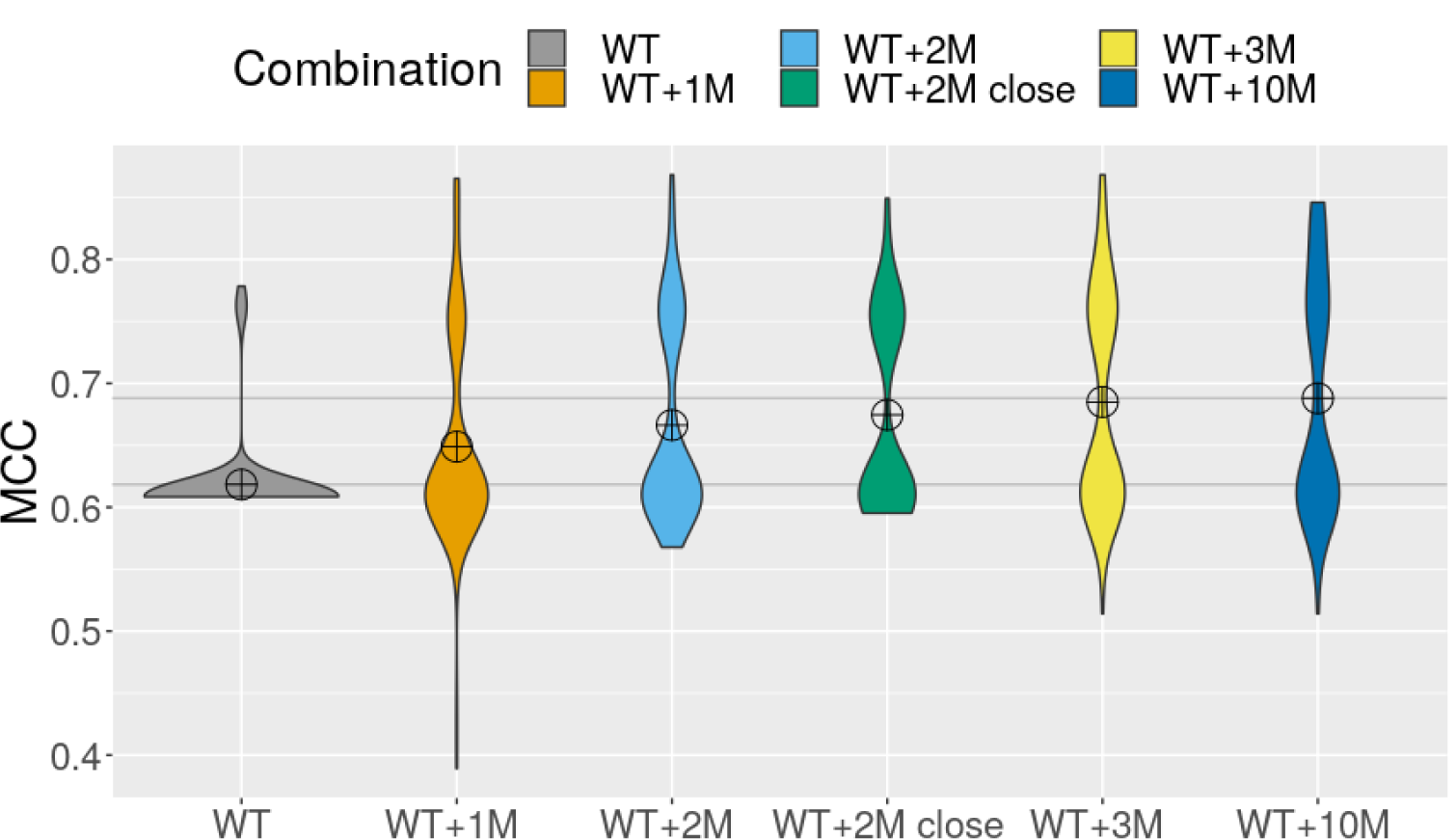
Distributions of predictive accuracies (MCC) of structures predicted by `IPANEMAP`, informed by Mutate-and-Map data, for DiLCrz (Cheng dataset (Cheng et al., 2017)). Predictions were repeated 100 times, informed both by the wild-type sequence and reactivity profile (WT), and with randomly-selected reactivity profiles associated with 1, 2, 3, or 10 single-point mutants (WT+*k*M). Pairs of mutants restricted to the 20 most similar to the WT were also considered (WT+2M close).

For the WT alone, the predictions of `IPANEMAP` were associated with 61.8% MCC, comparable to the worst condition of our previous dataset. Our predictions were generally stable and, in 94/100 runs, resulted in a structure having MCC between 60.5% and 61%. The remaining 6 runs showed improved MCC values, ranging from 76% to 78% and suggesting the existence of an alternative conformation in the pseudo-Boltzmann ensemble.

When the WT and a single random mutant (WT+1M) were jointly considered, the dispersion of the MCCs increased, and values ranging from 38.9% to 85% were observed.The average MCC in the presence of a single mutant increased to 64.9%,suggesting a significant positive contribution from the additional mutant profile (P-value= 1 *×* 10^−3^).

This trend is confirmed when two random mutants were considered in addition to the WT (WT+2M), with an average MCC across runs increasing to 66.6%. Interestingly, the dispersion of the MCC was lower for WT+2M than for WT+1M, with lower MCC values above 57%. This results suggest adding a second mutant profile has a role of *tie-breaker*, allowing to mitigate the deleterious effect of some outlier. However, outliers still played a disruptive role, and restricting chosen mutants to the 20 reactivity profiles most similar to the WT further improved the average MCC 2.8% to 67.7%, a significant improvement compared to WT+1M (P-value= 2.4 *×* 10^−2^).

The average MCC again increased in the presence of 3 mutants, to reach an average MCC of 69.3%, a substantial improvement that nevertheless fails to reach the level of statistical significance (P-value= 11 *×* 10^−2^). For 10 mutants, the average MCC and overall distribution remains highly similar to that obtained by considering three mutants, suggesting a plateau in the performances. Overall, these results support a notion of complementarity between the probing data produced across reasonably similar mutants, leading to a gradual decrease of the signal to noise ratio. The bimodal nature of the distribution also suggests the existence of two conformations, equally supported by the WT and the mutant profiles.

## Conclusion and discussion

We introduce `IPANEMAP`, an integrative method for the prediction of secondary structure models from multiple probing profiles. Based on the simple premise that good structural models should be thermodynamically stable and supported by most probing experiments, it uses a combination of stochastic backtrack in the pseudo-Boltzmann ensemble, coupled with a structural clustering to elect dominant conformations. This strategy is fast, generally stable, and leads to predictions that clearly benefit from the availability of multiple probing profiles, as demonstrated by detailed analyses performed of four datasets.

Our analyses reveal that integrating multiple profiles of at least three conditions allows to mitigate the effect of outlier conditions and, at times, to produce better secondary structure models than the one inferred only from the most accurate profile. A comparison of the joint performances to the average appears fair since no single reagent appears to systematically dominate in term of predictive capacity. It follows that a modeler cannot objectively favor *a priori* a probing condition/reagent (Yu et al., 2018). In other words, multiple experimental probing profiles, even combined in a pairwise fashion, can be used jointly to mitigate the empirical risk of a misprediction.

To the best of our knowledge, `IPANEMAP` currently represents the only available method for a joint automated consideration of multiple probing profiles. While hard constraints (Mathews et al., 2004) could in principle support multiple sources of probing data, their derivation from reactivities usually require a careful choice of cutoffs, making their automation virtually impossible. Overly liberal cutoffs will often induce contradictory constraints (i.e. no compatible structure) across conditions, while conservative cutoffs will lead to sparse constraints, practically wasting most of the probing-derived information. In fact the tediousness, and suboptimality, of the manual determination of those cutoffs while modeling HIV-1 structural elements (Deforges et al., 2017), was one of the main motivation behind the present work.

Current limitations of `IPANEMAP` include the lack of an explicit support for pseudoknots, which will be addressed in a future version, using polynomial-time dynamic programming (Janssen and Giegerich, 2015), or a iterative heuristic alternative (Hajdin et al., 2013). The inherently stochastic nature of the sampling scheme may also be challenging for certain types of analysis. However, despite its stochastic foundations, `IPANEMAP` is typically stable in its predictions, as can be seen in Figure 9 for all probing datasets produced over our DiLCrz case study. Interestingly, the most notable exception to the general stability is observed for the probing-free prediction, consistent with the widespread interpretation of SHAPE-induced pseudo-potentials as *focusing* the predictions onto a subset of the Boltzmann ensemble. A similar behavior is observed for the other datasets considered in our study (see Supp. Tables S2 and S5). Finally, the current clustering method induce a growth of time and memory requirements that scales quadratically with the number of samples. While this allows a joint analysis of a dozen of reactivity profiles in less than an hour on a personal computer, we expect the ability to push the number of samples will lead to better, even more reproducible, predictions. To that purpose, we will explore embedding techniques and linear-time community detection algorithms as alternatives to MBkM (Sculley, 2010).

**Figure 9:**
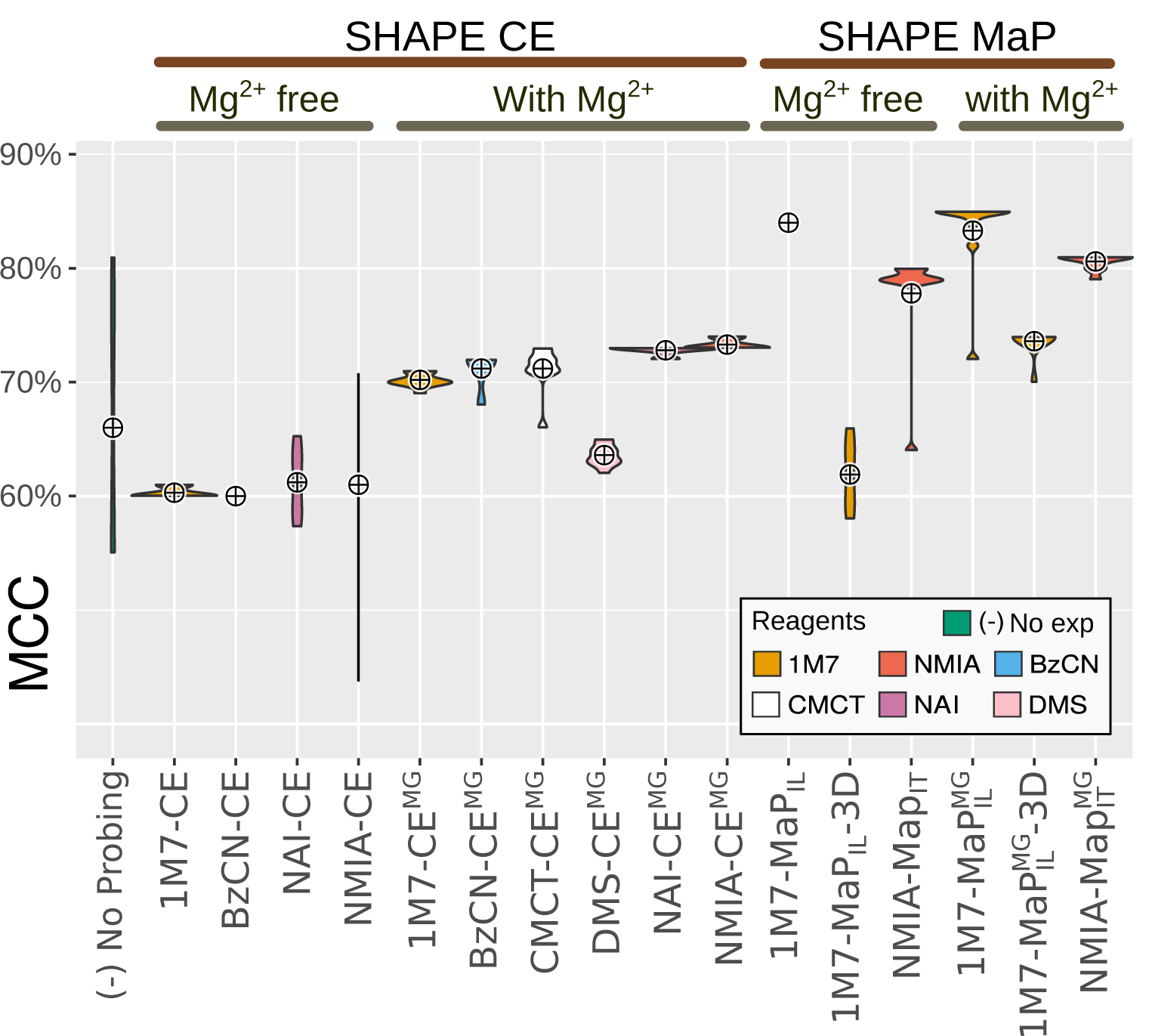
Stochastic variability of `IPANEMAP` predictions for DiLCrz dataset. MCC of predicted structures over 10 runs of `IPANEMAP`, in presence/absence of Mg^2+^, using both SHAPE CE and MaP protocols for a collection of reagents.

Future extensions also include addressing the dilemma of whether or not to include Mg^2+^ ions during the probing, in the perspective of a downstream computational prediction. On the one hand, RNA reaches its native 3D structure in the presence of Mg^2+^ ions (Knapp, 1989). However, such native structures include non WC pairings, and complex topological motifs such as pseudoknots, kissing loops, tetraloop interactions, that classic structure prediction algorithms do not explicitly capture. In contrast, in the presence of only monovalent ions, RNA is expected to form its helical domains, but leave its tertiary contacts unstable, with the notable exception of the G quadruplex. Therefore, probing the RNA without Mg^2+^ could provide a signal that is less deceptive, and thus more informative, to predicting algorithms. Unfortunately, destabilizing tertiary contacts increase the probability of misfolding, or the coexistence in solution of multiple conformations. In this work we probed DiLCrz, which contains many tertiary motifs, in the presence and absence of Mg^2+^. Remarkably, the most accurate predictions were obtained with 1M7 and NMIA irrespectively of the presence/absence of Mg^2+^, and despite the fact that reactivity profiles substantially diverged between the two conditions (see Supp. Figure S4). However, for other reagents/protocols, predictions appeared positively impacted (+10% average MCC) by the presence of Mg^2+^, painting a mixed picture that would deserve future investigations.

## Authors contributions

AS, BS, and YP designed the computational method. AS and YP implemented the computational method. DA and BS planned and carried out all probing experiments. AS and YP planned and carried out the computational experiments. AS, DA, BS and YP contributed to the interpretation of the results. All authors provided critical feedback and helped shape the research, analysis and manuscript

## Acknowledgements

The authors wish to express their gratitude to: Ronny Lorenz for developing a probing-informed version of RNAsubopt, allowing to sample the pseudo-Boltzmann ensemble; Nathalie Chamond for helpful suggestions; Benoit Masquida for the gift of DiLCrz plasmid and helpful discussions about protocols and DiLCrz structure; and an anonymous reviewer for helping us detect an error in our execution of `RSample`.

## Bibliography

C Brunel and P Romby. Probing RNA structure and RNA-ligand complexes with chemical probes. Methods Enzymol, 318:3–21, 2000. ISSN 0076-6879.

Steven Busan and Kevin M. Weeks. Accurate detection of chemical modifications in rna by mutational profiling (map) with shapemapper 2. RNA, 24(2):143–148, 2018. doi: 10.1261/rna.061945.117. URL http://rnajournal.cshlp.org/content/24/2/143.abstract.

Steven Busan, Chase A Weidmann, Arnab Sengupta, and Kevin M Weeks. Guidelines for SHAPE Reagent Choice and Detection Strategy for RNA Structure Probing Studies. Biochemistry (Mosc), 58:2655–2664, 2019. ISSN 1520-4995. doi: 10.1021/acs.biochem.8b01218.

S E Butcher and J M Burke. Structure-mapping of the hairpin ribozyme. Magnesium-dependent folding and evidence for tertiary interactions within the ribozyme-substrate complex. J Mol Biol, 244:52–63, 1994. ISSN 0022-2836. doi: 10.1006/jmbi.1994.1703.

Clarence Y Cheng, Wipapat Kladwang, Joseph D Yesselman, and Rhiju Das. RNA structure inference through chemical mapping after accidental or intentional mutations. Proc Natl Acad Sci U S A, 114(37):9876–9881, 2017. ISSN 1091-6490. doi: 10.1073/pnas.1619897114. URL http://www.ncbi.nlm.nih.gov/pubmed/28851837http://www.pubmedcentral.nih.gov/articlerender.fcgi?artid=PMC5603990.

Pablo Cordero and Rhiju Das. Rich RNA Structure Landscapes Revealed by Mutate-and-Map Analysis. PLoS Comput Biol, 11:e1004473, 2015. ISSN 1553-7358. doi: 10.1371/journal.pcbi.1004473.

Pablo Cordero, Wipapat Kladwang, Christopher C Vanlang, and Rhiju Das. Quantitative Dimethyl Sulfate Mapping for Automated RNA Secondary Structure Inference. Biochemistry (Mosc), 51(36):7037–7039, 2012a. doi: 10.1021/bi3008802. URL https://pubs.acs.org/doi/pdf/10.1021/bi3008802.

Pablo Cordero, Julius B. Lucks, and Rhiju Das. An RNA mapping data base for curating RNA structure mapping experiments. Bioinformatics, 28(22):3006–3008, 2012b. ISSN 13674803. doi: 10.1093/bioinformatics/bts554.

Jules Deforges, Sylvain de Breyne, Melissa Ameur, Nathalie Ulryck, Nathalie Chamond, Afaf Saaidi, Yann Ponty, Theophile Ohlmann, and Bruno Sargueil. Two ribosome recruitment sites direct multiple translation events within HIV1 Gag open reading frame. Nucleic Acids Res, 45:7382–7400, 2017. ISSN 1362-4962. doi: 10.1093/nar/gkx303.

Katherine E. Deigan, Tian W. Li, David H. Mathews, and Kevin M. Weeks. Accurate SHAPE-directed RNA structure determination. Proc Natl Acad Sci U S A, 106(1):97–102, 2009. ISSN 0027-8424. doi: 10.1073/pnas.0806929106. URL https://www.pnas.org/content/106/1/97.

Fei Deng, Mirko Ledda, Sana Vaziri, and Sharon Aviran. Data-directed RNA secondary structure prediction using probabilistic modeling. RNA, 22(8):1109–1119, 2016. ISSN 14699001. doi: 10.1261/rna.055756.115.

Y. Ding and Charles E. Lawrence. A statistical sampling algorithm for RNA secondary structure prediction. Nucleic Acids Res, 31(24):7280–7301, 2003. ISSN 1362-4962. doi: 10.1093/nar/gkg938. URL http://nar.oxfordjournals.org/content/31/24/7280.long.

C Ehresmann, F Baudin, M Mougel, P Romby, J P Ebel, and B Ehresmann. Probing the structure of RNAs in solution. Nucleic Acids Res, 15:9109–9128, 1987. ISSN 0305-1048. doi: 10.1093/nar/15.22.9109.

E Frezza, A Courban, A Allouche, B Sargueil, and S Pasquali. The Interplay between Molecular Flexibility and RNA Chemical Probing Reactivities Analyzed at the Nucleotide Level via an Extensive Molecular Dynamics Study. Methods, 162–163:108–127, 2019. doi: 10.1016/j.ymeth.2019.05.021.

Costin M Gherghe, Zahra Shajani, Kevin A Wilkinson, Gabriele Varani, and Kevin M Weeks. Strong correlation between SHAPE chemistry and the generalized NMR order parameter (S2) in RNA. J Am Chem Soc, 130: 12244–12245, 2008. ISSN 1520-5126. doi: 10.1021/ja804541s.

J Gorodkin, S L Stricklin, and G D Stormo. Discovering common stem-loop motifs in unaligned RNA sequences. Nucleic Acids Res, 29:2135–2144, 2001. ISSN 1362-4962.

Lauriane Gross, Quentin Vicens, Evelyne Einhorn, Audrey Noireterre, Laure Schaeffer, Lauriane Kuhn, Jean-Luc Imler, Gilbert Eriani, Carine Meignin, and Franck Martin. The IRES 5’UTR of the dicistrovirus cricket paralysis virus is a type III IRES containing an essential pseudoknot structure. Nucleic Acids Res, 45:8993–9004, 2017. ISSN 1362-4962. doi: 10.1093/nar/gkx622.

C. E. Hajdin, S. Bellaousov, W. Huggins, C. W. Leonard, D. H. Mathews, and K. M. Weeks. Accurate SHAPE-directed RNA secondary structure modeling, including pseudoknots. Proc Natl Acad Sci U S A, 110(14):5498–5503, 2013. ISSN 0027-8424. doi: 10.1073/pnas.1219988110.

Cécile H Herbreteau, Laure Weill, Didier Décimo, Déborah Prévôt, Jean-Luc Darlix, Bruno Sargueil, and Théophile Ohlmann. HIV-2 genomic RNA contains a novel type of IRES located downstream of its initiation codon. Nature structural & molecular biology, 12:1001–1007, 2005. ISSN 1545-9993. doi: 10.1038/nsmb1011.

Travis Hurst, Xiaojun Xu, Peinan Zhao, and Shi-Jie Chen. Quantitative Understanding of SHAPE Mechanism from RNA Structure and Dynamics Analysis. The journal of physical chemistry. B, 122:4771–4783, 2018. ISSN 1520-5207. doi: 10.1021/acs.jpcb.8b00575.

Laurie James and Bruno Sargueil. RNA secondary structure of the feline immunodeficiency virus 5’UTR and Gag coding region. Nucleic Acids Res, 36:4653–4666, 2008. ISSN 1362-4962. doi: 10.1093/nar/gkn447.

Stefan Janssen and Robert Giegerich. The rna shapes studio. Bioinformatics (Oxford, England), 31:423–425, February 2015. ISSN 1367-4811. doi: 10.1093/bioinformatics/btu649.

Fethullah Karabiber, Jennifer L McGinnis, Oleg V Favorov, and Kevin M Weeks. QuShape: rapid, accurate, and best-practices quantification of nucleic acid probing information, resolved by capillary electrophoresis. RNA, 19:63–73, 2013. ISSN 1469-9001. doi: 10.1261/rna.036327.112.

G Knapp. Enzymatic approaches to probing of RNA secondary and tertiary structure. Methods Enzymol, 180: 192–212, 1989. ISSN 0076-6879.

Christopher A Lavender, Ronny Lorenz, Ge Zhang, Rita Tamayo, Ivo L Hofacker, and Kevin M Weeks. Model-Free RNA Sequence and Structure Alignment Informed by SHAPE Probing Reveals a Conserved Alternate Secondary Structure for 16S rRNA. PLoS Comput Biol, 11:e1004126, 2015. ISSN 1553-7358. doi: 10.1371/journal.pcbi.1004126.

Ronny Lorenz, Stephan H Bernhart, Christian Höner Zu Siederdissen, Hakim Tafer, Christoph Flamm, Peter F Stadler, and Ivo L Hofacker. ViennaRNA Package 2.0. Algorithms for molecular biology : AMB, 6:26, 2011. ISSN 1748-7188. doi: 10.1186/1748-7188-6-26.

Ronny Lorenz, Dominik Luntzer, Ivo L Hofacker, Peter F Stadler, and Michael T Wolfinger. SHAPE directed RNA folding. Bioinformatics, 32:145–147, 2016. ISSN 1367-4811. doi: 10.1093/bioinformatics/btv523.

Xiang Jun Lu, Harmen J. Bussemaker, and Wilma K. Olson. DSSR: An integrated software tool for dissecting the spatial structure of RNA. Nucleic Acids Res, 43(21):e142, 2015. ISSN 13624962. doi: 10.1093/nar/gkv716.

Zhi John Lu, Jason W Gloor, and David H Mathews. Improved RNA secondary structure prediction by maximizing expected pair accuracy. RNA, 15(10):1805–1813, 2009.

David H Mathews, Matthew D Disney, Jessica L Childs, Susan J Schroeder, Michael Zuker, and Douglas H Turner. Incorporating chemical modification constraints into a dynamic programming algorithm for prediction of RNA secondary structure. Proc Natl Acad Sci U S A, 101(19):7287–92, 2004. URL http://www.pnas.org/content/101/19/7287.full.

Christopher A. Mattson and Achille Messac. Pareto frontier based concept selection under uncertainty, with visualization. Optimization and Engineering, 6(1):85–115, 2005.

J S McCaskill. The equilibrium partition function and base pair binding probabilities for RNA secondary structure. Biopolymers, 29(6-7):1105–1119, 1990.

Jennifer L McGinnis, Jack A Dunkle, Jamie H D Cate, and Kevin M Weeks. The mechanisms of RNA SHAPE chemistry. J Am Chem Soc, 134:6617–6624, 2012. ISSN 1520-5126. doi: 10.1021/ja2104075.

M. Meyer, H. Nielsen, V. Olieric, P. Roblin, S. D. Johansen, E. Westhof, and B. Masquida. Speciation of a group I intron into a lariat capping ribozyme. Proc. Natl. Acad. Sci. U.S.A., 111(21):7659–7664, 2014. ISSN 0027-8424. doi: 10.1073/pnas.1322248111.

Zhichao Miao, Ryszard W Adamiak, Maciej Antczak, Robert T Batey, Alexander J Becka, Marcin Biesiada, Michał J Boniecki, Janusz M Bujnicki, Shi-Jie Chen, Clarence Yu Cheng, Fang-Chieh Chou, Adrian R Ferré-D’Amaré, Rhiju Das, Wayne K Dawson, Feng Ding, Nikolay V Dokholyan, Stanisław Dunin-Horkawicz, Caleb Geniesse, Kalli Kappel, Wipapat Kladwang, Andrey Krokhotin, Grzegorz E Łach, François Major, Thomas H Mann, Marcin Magnus, Katarzyna Pachulska-Wieczorek, Dinshaw J Patel, Joseph A Piccirilli, Mariusz Popenda, Katarzyna J Purzycka, Aiming Ren, Greggory M Rice, John Santalucia, Joanna Sarzynska, Marta Szachniuk, Arpit Tandon, Jeremiah J Trausch, Siqi Tian, Jian Wang, Kevin M Weeks, Benfeard Williams, Yi Xiao, Xiaojun Xu, Dong Zhang, Tomasz Zok, and Eric Westhof. RNA-Puzzles Round III: 3D RNA structure prediction of five riboswitches and one ribozyme. RNA, 23:655–672, 2017. ISSN 1469-9001. doi: 10.1261/rna.060368.116.

Vojtěch Mlýnský and Giovanni Bussi. Molecular Dynamics Simulations Reveal an Interplay between SHAPE Reagent Binding and RNA Flexibility. The journal of physical chemistry letters, 9:313–318, 2018. ISSN 1948-7185. doi: 10.1021/acs.jpclett.7b02921.

D Moazed, S Stern, and H F Noller. Rapid chemical probing of conformation in 16 S ribosomal RNA and 30 S ribosomal subunits using primer extension. J Mol Biol, 187:399–416, 1986. ISSN 0022-2836.

Jean-Christophe Paillart, Markus Dettenhofer, Xiao-fang Yu, Chantal Ehresmann, Bernard Ehresmann, and Roland Marquet. First snapshots of the hiv-1 rna structure in infected cells and in virions. Journal of Biological Chemistry, 279(46):48397–48403, 2004. doi: 10.1074/jbc.M408294200. URL http://www.jbc.org/content/279/46/48397.abstract.

Fabian Pedregosa, Gaël Varoquaux, Alexandre Gramfort, Vincent Michel, Bertrand Thirion, Olivier Grisel, Mathieu Blondel, Peter Prettenhofer, Ron Weiss, Vincent Dubourg, Jake Vanderplas, Alexandre Passos, David Cournapeau, Matthieu Brucher, Matthieu Perrot, and Édouard Duchesnay. Scikit-learn: Machine Learning in Python. Journal of Machine Learning Research, 12:2825–2830, 2012. URL http://dl.acm.org/citation.cfm?id=2078195{%}5Cnhttp://arxiv.org/abs/1201.0490.

Jessica S. Reuter and David H. Mathews. RNAstructure: Software for RNA secondary structure prediction and analysis. BMC Bioinf, 11:129, 2010. ISSN 14712105. doi: 10.1186/1471-2105-11-129.

Greggory M Rice, Christopher W Leonard, and Kevin M Weeks. RNA secondary structure modeling at consistent high accuracy using differential SHAPE. RNA, 20:846–854, 2014. ISSN 1469-9001. doi: 10.1261/rna.043323.113.

P J Romaniuk, I L de Stevenson, C Ehresmann, P Romby, and B Ehresmann. A comparison of the solution structures and conformational properties of the somatic and oocyte 5S rRNAs of Xenopus laevis. Nucleic Acids Res, 16:2295–2312, 1988. ISSN 0305-1048. doi: 10.1093/nar/16.5.2295.

D Sculley. Web-scale k-means clustering. In Proceedings of the 19th international conference on World Wide Web (WWW’10), pages 1177–1178, 2010. URL http://portal.acm.org/citation.cfm?doid=1772690.1772862.

Alec N Sexton, Peter Y Wang, Michael Rutenberg-Schoenberg, and Matthew D Simon. Interpreting Reverse Transcriptase Termination and Mutation Events for Greater Insight into the Chemical Probing of RNA. Biochemistry (Mosc), 56:4713–4721, 2017. ISSN 1520-4995. doi: 10.1021/acs.biochem.7b00323.

Sandra Smit, Kristian Rother, Jaap Heringa, and Rob Knight. From knotted to nested RNA structures: a variety of computational methods for pseudoknot removal. RNA, 14:410–416, 2008. ISSN 1469-9001. doi: 10.1261/rna.881308.

Matthew J Smola, Greggory M Rice, Steven Busan, Nathan A Siegfried, and Kevin M Weeks. Selective 2’-hydroxyl acylation analyzed by primer extension and mutational profiling (SHAPE-MaP) for direct, versatile and accurate RNA structure analysis. Nat Protoc, 10:1643–1669, 2015. ISSN 1750-2799. doi: 10.1038/nprot.2015.103.

Srinivas Somarowthu, Michal Legiewicz, Isabel Chillón, Marco Marcia, Fei Liu, and Anna Marie Pyle. HOTAIR forms an intricate and modular secondary structure. Mol Cell, 58:353–361, 2015. ISSN 1097-4164. doi: 10.1016/j.molcel.2015.03.006.

A. Spasic, S. M. Assmann, P. C. Bevilacqua, and D. H. Mathews. Modeling RNA secondary structure folding ensembles using SHAPE mapping data. Nucleic Acids Res, 46(1):314–323, 2017. doi: 10.1093/nar/gkx1057. URL https://www.ncbi.nlm.nih.gov/pubmed/29177466.

Kady-Ann Steen, Greggory M Rice, and Kevin M Weeks. Fingerprinting noncanonical and tertiary RNA structures by differential SHAPE reactivity. J Am Chem Soc, 134:13160–13163, 2012. ISSN 1520-5126. doi: 10.1021/ja304027m.

Douglas H Turner and David H Mathews. NNDB: the nearest neighbor parameter database for predicting stability of nucleic acid secondary structure. Nucleic Acids Res, 38:D280–D282, 2010. ISSN 1362-4962. doi: 10.1093/nar/gkp892.

Stefan Washietl, Ivo L. Hofacker, Peter F. Stadler, and Manolis Kellis. RNA folding with soft constraints: Reconciliation of probing data and thermodynamic secondary structure prediction. Nucleic Acids Res, 40(10): 4261–4272, 2012. ISSN 03051048. doi: 10.1093/nar/gks009.

Laure Weill, Dominique Louis, and Bruno Sargueil. Selection and evolution of NTP-specific aptamers. Nucleic Acids Res, 32:5045–5058, 2004. ISSN 1362-4962. doi: 10.1093/nar/gkh835.

Kevin A Wilkinson, Edward J Merino, and Kevin M Weeks. Selective 2’-hydroxyl acylation analyzed by primer extension (SHAPE): quantitative RNA structure analysis at single nucleotide resolution. Nat Protoc, 1:1610–1616, 2006. ISSN 1750-2799. doi: 10.1038/nprot.2006.249.

Yang Wu, Binbin Shi, Xinqiang Ding, Tong Liu, Xihao Hu, Kevin Y. Yip, Zheng Rong Yang, David H. Mathews, and Zhi John Lu. Improved prediction of RNA secondary structure by integrating the free energy model with restraints derived from experimental probing data. Nucleic Acids Res, 43(15):7247–7259, 2015. ISSN 13624962. doi: 10.1093/nar/gkv706.

Zhenjiang Xu, Anthony Almudevar, and David H Mathews. Statistical evaluation of improvement in RNA secondary structure prediction. Nucleic acids research, 40:e26, February 2012. ISSN 1362-4962. doi: 10.1093/nar/gkr1081.

Angela M Yu, Molly E. Evans, and Julius B. Lucks. Estimating RNA structure chemical probing reactivities from reverse transcriptase stops and mutations. BioRxiv 292532, 2018. URL https://www.biorxiv.org/content/early/2018/03/30/292532.

Kourosh Zarringhalam, Michelle M. Meyer, Ivan Dotu, Jeffrey H. Chuang, and Peter Clote. Integrating Chemical Footprinting Data into RNA Secondary Structure Prediction. PLoS One, 7(10):e45160, 2012. ISSN 19326203. doi: 10.1371/journal.pone.0045160.

A J Zaug and T R Cech. Analysis of the structure of tetrahymena nuclear rnas in vivo: telomerase rna, the self-splicing rrna intron, and u2 snrna. RNA, 1(4):363–74, 1995. URL http://rnajournal.cshlp.org/content/1/4/363.abstract.

