## Supplementary material for "IPANEMAP: Integrative Probing Analysis of Nucleic Acids Empowered by Multiple Accessibility Profiles"

### 1 Multiprobing benchmark (Cordero *et al* [1] dataset)

#### 1.1 GM values of individual predictions

| | `IPANEMAP` | | | | | | | | `RNAfold` – MFE mode | | | | `RNAfold` – MEA mode | | | |
|---|---|---|---|---|---|---|---|---|---|---|---|---|---|---|---|---|
| **CMCT** | ○ | ● | ○ | ○ | ○ | ● | ● | ● | ○ | ○ | ● | ○ | ○ | ○ | ● | ○ |
| **NMIA** | ○ | ○ | ● | ○ | ● | ○ | ● | ● | ○ | ○ | ○ | ● | ○ | ○ | ○ | ● |
| **DMS** | ○ | ○ | ○ | ● | ● | ● | ○ | ● | ○ | ● | ○ | ○ | ○ | ● | ○ | ○ |
| 5SRNA,E.Coli | 0.25 | 0.238 | 0.247 | 0.244 | 0.247 | 0.247 | 0.254 | 0.241 | 0.241 | 0.686 | 0.254 | 0.686 | 0.241 | 0.686 | 0.269 | 0.686 |
| glycineriboswitch,F.nucleatum | 0.568 | 0.627 | 0.868 | 0.952 | 0.868 | 0.658 | 0.868 | 0.868 | 0.306 | 0.313 | 0.395 | 0.313 | 0.593 | 0.593 | 0.693 | 0.6 |
| cidGMPriboswitch,V.Cholerae | 0.654 | 0.77 | 0.654 | 0.77 | 0.654 | 0.77 | 0.654 | 0.77 | 0.77 | 0.77 | 0.667 | 0.77 | 0.77 | 0.72 | 0.667 | 0.77 |
| P4P6domain | 0.864 | 0.856 | 0.881 | 0.864 | 0.864 | 0.856 | 0.864 | 0.856 | 0.837 | 0.808 | 0.773 | 0.714 | 0.845 | 0.808 | 0.781 | 0.722 |
| adenineriboswitch,add | 1 | 0.956 | 1 | 0.977 | 1 | 0.956 | 1 | 1 | 1 | 0.333 | 0.41 | 1 | 1 | 0.333 | 0.356 | 0.306 |
| tRNAphenylalanineyeast | 0.286 | 0.746 | 0.976 | 0.746 | 0.976 | 0.746 | 0.976 | 0.976 | 0.976 | 0.976 | 0.976 | 0.976 | 0.334 | 0.976 | 0.976 | 0.976 |
| Average | 0.604 | 0.7 | 0.77 | 0.76 | 0.77 | 0.705 | 0.77 | 0.785 | 0.69 | 0.65 | 0.58 | 0.74 | 0.63 | 0.69 | 0.62 | 0.68 |

**Table S1:** GM of the structures predicted with **`IPANEMAP`**, compared to predictions of **`RNAfold`** in energy mimization (MFE) and accuracy maximization (MEA) modes.

#### 1.2 Stablity analysis

| | NMIA | | | DMS | | | CMCT | | | NMIA+DMS | | | NMIA+CMCT | | | DMS+CMCT | | | NMIA+DMS+CMCT | | |
|---|---|---|---|---|---|---|---|---|---|---|---|---|---|---|---|---|---|---|---|---|---|
| Replicates | Run 1 | Run 2 | Run 3 | Run 1 | Run 2 | Run 3 | Run 1 | Run 2 | Run 3 | Run 1 | Run 2 | Run 3 | Run 1 | Run 2 | Run 3 | Run 1 | Run 2 | Run 3 | Run 1 | Run 2 | Run 3 |
| 5SRNA,E.Coli | 0.247 | 0.25 | 0.247 | 0.254 | 0.244 | 0.244 | 0.238 | 0.238 | 0.238 | 0.247 | 0.244 | 0.247 | 0.257 | 0.238 | 0.238 | 0.247 | 0.247 | 0.244 | 0.241 | 0.241 | 0.241 |
| glycineriboswitch,F.nucleatum | 0.868 | 0.868 | 0.868 | 0.952 | 0.952 | 0.952 | 0.868 | 0.634 | 0.627 | 0.868 | 0.868 | 0.868 | 0.868 | 0.868 | 0.868 | 0.667 | 0.658 | 0.658 | 0.868 | 0.868 | 0.868 |
| cidGMPriboswitch,V.Cholerae | 0.631 | 0.631 | 0.631 | 0.77 | 0.77 | 0.77 | 0.667 | 0.77 | 0.77 | 0.631 | 0.631 | 0.756 | 0.77 | 0.77 | 0.77 | 0.77 | 0.77 | 0.77 | 0.77 | 0.77 | 0.77 |
| P4P6domain | 0.864 | 0.881 | 0.881 | 0.864 | 0.864 | 0.856 | 0.856 | 0.856 | 0.856 | 0.864 | 0.864 | 0.864 | 0.864 | 0.864 | 0.864 | 0.856 | 0.856 | 0.856 | 0.856 | 0.856 | 0.856 |
| adenineriboswitch,ad | 1 | 1 | 1 | 0.956 | 0.956 | 0.956 | 1 | 0.956 | 0.956 | 1 | 1 | 1 | 1 | 1 | 1 | 0.956 | 0.956 | 0.956 | 1 | 1 | 1 |
| tRNAphenylalanineyeast | 0.953 | 0.953 | 0.976 | 0.976 | 0.746 | 0.976 | 0.746 | 0.746 | 0.746 | 0.976 | 0.976 | 0.976 | 0.976 | 0.976 | 0.97 | 0.746 | 0.746 | 0.746 | 0.746 | 0.976 | 0.976 |

**Table S2:** GM of **`IPANEMAP`** predictions over three consecutive runs from up 3 sources of probing data.

### 2 Mono probing benchmark (Hajdin *et al* [2] dataset)

| RNA ID | Description | | |
|---|---|---|---|
| A | Pre-Q1 riboswitch, B. subtilis | M | SARS corona virus pseudoknot |
| B | 5' domain of 16S rRNA, E. coli | N | 5S rRNA, E. coli |
| C | Signal recogniton particle RNA human | O | cyclic-di-GMP riboswitch, V. cholerae |
| D | Group I intron, Azoarcus sp. | P | 5' domain of 23S rRNA, E. coli |
| E | HIV 15 prime pseudoknot domain | Q | RNase PB. subtilis |
| F | Telomerase pseudoknot human | R | Group I Intron, T. thermophila |
| G | SAM I riboswitch, T. tengcongensis | S | Hepatitis C virus IRES domain |
| H | tRNA(phe), E. coli | T | 5' domain of 16S rRNA, H. volcanii |
| I | P546 domain, bI3 group I intron | U | Group II intron, O. iheyensis |
| J | TPP riboswitch, E. coli | V | tRNA(asp), yeast |
| K | Adenine riboswitch, V. vulnificus | W | Lysine riboswitch, T. maritima |
| L | Fluoride riboswitch, P. syringae | X | M-Box riboswitch, B. subtilis |

**Table S3:** Labels used as shorthands for the 24 RNAs in the Hajdin *et al.* dataset [2].

#### 2.1 Detailed results of `IPANEMAP` and `Rsample`

| RNA | Length | RSample Sensitivity | PPV | GM | IPANEMAP Sensitivity | PPV | GM |
|---|---|---|---|---|---|---|---|
| A | 34 | 0.625 | 1.000 | 0.791 | 0.625 | 1 | 0.791 |
| B | 530 | 0.818 | 0.747 | 0.781 | 0.8986 | 0.8062 | 0.851 |
| C | 301 | 0.590 | 0.590 | 0.590 | 0.6 | 0.6 | 0.6 |
| D | 214 | 0.746 | 0.870 | 0.806 | 0.7302 | 0.8519 | 0.789 |
| E | 500 | 0.487 | 0.503 | 0.495 | 0.5263 | 0.5517 | 0.539 |
| F | 47 | 0.600 | 0.750 | 0.671 | 0.6 | 0.75 | 0.671 |
| G | 118 | 0.795 | 0.912 | 0.851 | 0.7692 | 0.8333 | 0.801 |
| H | 76 | 0.952 | 0.952 | 0.952 | 0.7619 | 0.7619 | 0.762 |
| I | 155 | 0.946 | 0.964 | 0.955 | 0.9821 | 0.9821 | 0.982 |
| J | 79 | 0.636 | 0.583 | 0.609 | 0.9545 | 0.875 | 0.914 |
| K | 71 | 1.000 | 1.000 | 1.000 | 1 | 0.9545 | 0.977 |
| L | 66 | 0.625 | 0.667 | 0.646 | 0.625 | 0.7143 | 0.668 |
| M | 79 | 0.654 | 0.654 | 0.654 | 0.6538 | 0.68 | 0.667 |
| N | 120 | 0.857 | 0.833 | 0.845 | 0.9714 | 0.9189 | 0.945 |
| O | 97 | 0.929 | 0.929 | 0.929 | 0.9286 | 0.8966 | 0.912 |
| P | 511 | 0.908 | 0.783 | 0.843 | 0.8908 | 0.7681 | 0.827 |
| Q | 401 | 0.670 | 0.726 | 0.697 | 0.7304 | 0.75 | 0.74 |
| R | 425 | 0.817 | 0.811 | 0.814 | 0.9084 | 0.8815 | 0.895 |
| S | 336 | 0.798 | 0.865 | 0.831 | 0.8077 | 0.866 | 0.836 |
| T | 473 | 0.889 | 0.815 | 0.851 | 0.8819 | 0.8194 | 0.85 |
| U | 412 | 0.485 | 0.533 | 0.508 | 0.75 | 0.8462 | 0.797 |
| V | 75 | 1.000 | 1.000 | 1.000 | 0.619 | 0.5652 | 0.591 |
| W | 174 | 0.810 | 0.895 | 0.851 | 0.8095 | 0.9808 | 0.891 |
| X | 154 | 0.875 | 0.913 | 0.894 | 0.875 | 0.913 | 0.894 |

**Table S4:** `IPANEMAP` prediction performance compared to `RSample` tool

#### 2.2 Stability analysis

| RNA | Length | Run1 | Run2 | Run3 | Run4 | Run5 | Run6 | Run7 | Run8 | Run9 | Run10 | Average | Std Dev | **MFE-RNAfold** | **MEA-RNAfold** |
|---|---|---|---|---|---|---|---|---|---|---|---|---|---|---|---|
| A | 34 | 0.791 | 0.791 | 0.791 | 0.791 | 0.791 | 0.791 | 0.791 | 0.791 | 0.791 | 0.722 | 0.7841 | 0.0207 | 0.791 | 0.791 |
| B | 530 | 0.851 | 0.851 | 0.851 | 0.851 | 0.851 | 0.851 | 0.851 | 0.851 | 0.851 | 0.851 | 0.851 | 0 | 0.851 | 0.849 |
| C | 301 | 0.6 | 0.6 | 0.6 | 0.6 | 0.6 | 0.6 | 0.6 | 0.6 | 0.6 | 0.6 | 0.6 | 0 | 0.6 | 0.597 |
| D | 214 | 0.789 | 0.789 | 0.789 | 0.781 | 0.781 | 0.781 | 0.781 | 0.781 | 0.781 | 0.781 | 0.7834 | 0.0037 | 0.768 | 0.761 |
| E | 500 | 0.539 | 0.539 | 0.539 | 0.539 | 0.539 | 0.539 | 0.539 | 0.535 | 0.541 | 0.515 | 0.5364 | 0.0073 | 0.562 | 0.524 |
| F | 47 | 0.671 | 0.671 | 0.671 | 0.671 | 0.671 | 0.671 | 0.671 | 0.671 | 0.671 | 0.671 | 0.671 | 0 | 0.671 | 0.671 |
| G | 118 | 0.801 | 0.801 | 0.801 | 0.801 | 0.801 | 0.801 | 0.801 | 0.801 | 0.801 | 0.801 | 0.801 | 0 | 0.892 | 0.879 |
| H | 76 | 1 | 1 | 1 | 1 | 1 | 1 | 1 | 0.762 | 0.762 | 0.781 | 0.9305 | 0.1063 | 1 | 1 |
| I | 155 | 0.982 | 0.982 | 0.982 | 0.982 | 0.982 | 0.982 | 0.982 | 0.982 | 0.982 | 0.964 | 0.9802 | 0.0054 | 0.964 | 0.982 |
| J | 79 | 0.914 | 0.914 | 0.914 | 0.914 | 0.914 | 0.914 | 0.914 | 0.914 | 0.597 | 0.889 | 0.8798 | 0.0946 | 0.914 | 0.914 |
| K | 71 | 1 | 1 | 1 | 1 | 1 | 1 | 1 | 0.977 | 0.977 | 0.977 | 0.9931 | 0.0105 | 1 | 1 |
| L | 66 | 0.668 | 0.668 | 0.668 | 0.668 | 0.668 | 0.668 | 0.668 | 0.668 | 0.668 | 0.668 | 0.668 | 0 | 0.668 | 0.668 |
| M | 79 | 0.706 | 0.706 | 0.706 | 0.706 | 0.706 | 0.706 | 0.721 | 0.721 | 0.667 | 0.745 | 0.709 | 0.0184 | 0.692 | 0.692 |
| N | 120 | 0.945 | 0.945 | 0.945 | 0.945 | 0.945 | 0.945 | 0.812 | 0.812 | 0.812 | 0.823 | 0.8929 | 0.064 | 0.945 | 0.914 |
| O | 97 | 0.912 | 0.912 | 0.912 | 0.912 | 0.912 | 0.912 | 0.912 | 0.912 | 0.929 | 0.929 | 0.9154 | 0.0068 | 0.948 | 0.948 |
| P | 511 | 0.824 | 0.824 | 0.824 | 0.821 | 0.821 | 0.818 | 0.818 | 0.833 | 0.83 | 0.827 | 0.824 | 0.0046 | 0.831 | 0.828 |
| Q | 401 | 0.74 | 0.74 | 0.74 | 0.74 | 0.74 | 0.74 | 0.74 | 0.74 | 0.74 | 0.74 | 0.74 | 0 | 0.813 | 0.75 |
| R | 425 | 0.895 | 0.895 | 0.895 | 0.895 | 0.895 | 0.895 | 0.895 | 0.895 | 0.895 | 0.895 | 0.895 | 0 | 0.885 | 0.92 |
| S | 336 | 0.836 | 0.836 | 0.836 | 0.836 | 0.836 | 0.836 | 0.836 | 0.836 | 0.836 | 0.831 | 0.8355 | 0.0015 | 0.831 | 0.831 |
| T | 473 | 0.85 | 0.85 | 0.85 | 0.85 | 0.85 | 0.85 | 0.85 | 0.85 | 0.85 | 0.85 | 0.85 | 0 | 0.863 | 0.863 |
| U | 412 | 0.929 | 0.929 | 0.929 | 0.929 | 0.81 | 0.81 | 0.81 | 0.806 | 0.806 | 0.925 | 0.8683 | 0.06 | 0.929 | 0.843 |
| V | 75 | 0.591 | 0.591 | 0.591 | 0.591 | 0.591 | 0.591 | 0.591 | 0.591 | 0.591 | 0.901 | 0.622 | 0.093 | 0.591 | 0.591 |
| W | 174 | 0.891 | 0.891 | 0.891 | 0.891 | 0.891 | 0.823 | 0.823 | 0.823 | 0.823 | 0.823 | 0.857 | 0.034 | 0.842 | 0.823 |
| X | 154 | 0.894 | 0.894 | 0.894 | 0.894 | 0.894 | 0.894 | 0.894 | 0.894 | 0.894 | 0.894 | 0.894 | 0 | 0.894 | 0.894 |

**Table S5:** MCC of predicted structures with `IPANEMAP` through 10 runs. The Average and the standard deviation (Sd) of the MCC over the 10 runs are reported alongside with the MCC of the MFE and MEA structures obtained with `RNAfold`.

.

### 3 Case-study: Lariat capping ribozyme of *Didymium iridis*

#### 3.1 Validation of accessibility profiles: Agreement with native and predicted structures

We assess and validate the overall quality of probing data by defining a natural notion of **agreement** between a reactivity profile and a structure. To that purpose, we consider a **proportion $\alpha$ of the most and least reactive positions**, initially considering $\alpha = 30\%$. Then, we investigate whether positions associated with extreme reactivity values are indeed paired or unpaired within the native structure.

**Agreement metrics.** More precisely, given a secondary structure $S$ of length $n$, we define the **1D projection** of $S$ as the vector $v_S$ such that

$$\forall i \in [1, n], v_S(i) = \begin{cases} 1 & \text{if } i \text{ unpaired in } S \\ 0 & \text{otherwise.} \end{cases}$$

Given a vector $r$ of reactivities, we define a **discrete reactivity vector** $d_r^\alpha$ as

$$\forall i \in [1, n], d_r^\alpha(i) = \begin{cases} 1 & \text{if } r_i \text{ within the } \alpha \text{ largest reactivity values,} \\ 0 & \text{if } r_i \text{ within the } \alpha \text{ lowest reactivity values,} \\ 1/2 & \text{otherwise (including if not available).} \end{cases}$$

By focusing on the highest and lowest values, we avoid the influence of outliers, and hope to partially mitigate the natural variability of probing reactivities.

We now define the **normalized divergence** $||r, S||$ of a reactivity profile $r$ from a structure $S$ as

$$||r, S|| = \frac{\sum_{i=1}^{n} |v_S(i) - d_r(i)| - (1 - 2\alpha) \times \frac{n}{2}}{n \times 2\alpha}.$$

A correcting term counterbalances the $1/2$ associated with medium values, leading to an overall divergence of $(1 - 2\alpha) \times \frac{n}{2}$ regardless of the quality of the profile. The divisor term of $n \times 2\alpha$ accounts for the fact that, after compensating for the contribution of the $[\alpha, 1 - \alpha]$ percentile, only a proportion $2\alpha$ of the original values is effectively covered by the sum. Implementing those two corrections leads to a final value scaling between $0$ (all highly reactive pos. unpaired, all poorly reactive pos. paired) and $1$ (all highly reactive pos. paired, all poorly reactive pos. unpaired).

Finally, we simply define the **agreement of a reactivity profile** $r$ **with a structure** $S$ as

$$\text{Agreement}(r, S) = 1 - ||r, S||.$$

**Results.** We computed, and report in Table 1, the agreement of our 16 reactivity profiles with both the native and `IPANEMAP` prediction when considering the $\alpha$ percentile of highest/lowest reactivities ($\alpha = 30\%$). A cursory inspection of the table reveals a **good overall agreement of the measured reactivities with the native structure**, involving $61\%$ of the positions on average. This agreement increases to an average $70\%$ when $\alpha$ is set to $15\%$, indicating a good correlation between raw reactivities values and paired/unpaired status in the native structure. Also, expectedly, the agreement to the native structure appears to be positively correlated with the MCC induced by the reactivity profile. However, this relationship is not strict, as revealed by the $60\%$ agreement to the native observed for 1M7-MAP$_{\text{IL}}$. While superficially surprising, this observation is consistent with the notion that probing data do not need to inform on many different positions to improve the quality of predictions.

We tested the **statistical significance** of our observation by computing the agreement distribution in the absence of structural information. To that purpose, we generated random reactivity profiles, drawing a random reactivity uniformly in $[0, 1]$ for each position. The resulting distribution can be perfectly fitted to a Normal distribution with mean agreement $\mu = 49.96\%$ and standard deviation $\sigma = 4.51\%$.

| | | | Agreement$(R, S)$ (%) | | |
|---|---|---|---|---|---|
| Num | Cluster | Condition $R$ | $S := \mathcal{N}$ | $S := \mathcal{P}$ | MCC% |
| 1 | C | 1M7-MAP$^{MG}_{IL}$ | 70 | 77 | 85 |
| 2 | A | 1M7-MAP$_{IL}$ | 60 | 68 | 84 |
| 3 | A | NMIA-MAP$^{MG}_{IT}$ | 73 | 74 | 81 |
| 4 | A | NMIA-MAP$_{IT}$ | 62 | 70 | 80 |
| 5 | C | 1M7-MAP$^{MG}_{IL}$-3D | 62 | 58 | 74 |
| 6 | E | NMIA-CE$^{MG}$ | 61 | 54 | 73 |
| 7 | G | NAI-CE$^{MG}$ | 61 | 62 | 73 |
| 8 | H | BzCN-CE$^{MG}$ | 70 | 62 | 71 |
| 9 | E | 1M7-CE$^{MG}$ | 58 | 52 | 70 |
| 10 | G | CMCT-CE$^{MG}$ | 55 | 51 | 70 |
| 11 | D | 1M7-MAP$_{IL}$-3D | 61 | 67 | 64 |
| 12 | F | DMS-CE$^{MG}$ | 62 | 62 | 63 |
| 13 | B | NMIA-CE | 62 | 69 | 61 |
| 14 | B | 1M7-CE | 59 | 62 | 60 |
| 15 | B | BzCN-CE | 54 | 49 | 60 |
| 16 | B | NAI-CE | 56 | 59 | 60 |

**Figure S1:** Analysis of agreements for reactivity profiles generated in various experimental conditions with both native ($\mathcal{N}$) and **IPANEMAP**-predicted ($\mathcal{P}$) structure.

**All reactivity profiles showed better agreement with the native structure than a random reactivity profile**, by a margin of **up to five standard deviations**, and always at least one standard deviation, better than the expected value of random profiles. This leads to **significantly enriched agreements** (P-values(Agreement)$<$5%) for 13 out of our 16 reactivity profiles, the remaining outliers having associated P-values of 18.5%, 13.2% and 9% respectively. Overall, these observations support the notion of informative probing experiments.

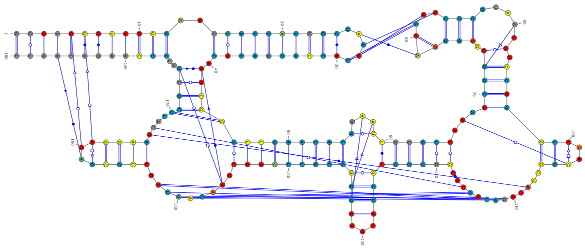

1 – 1M7-MaP$_{\text{IL}}^{\text{MG}}$ (Cluster C ●)

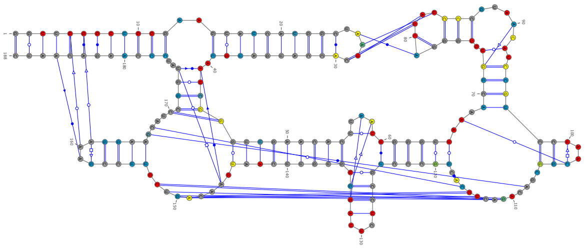

2 – 1M7-MaP$_{\text{IL}}$ (Cluster A ●)

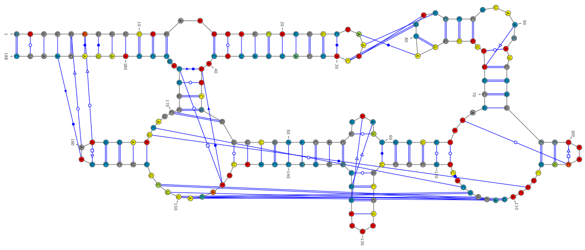

3 – NMIA-MaP$_{\text{IT}}^{\text{MG}}$ (Cluster A ●)

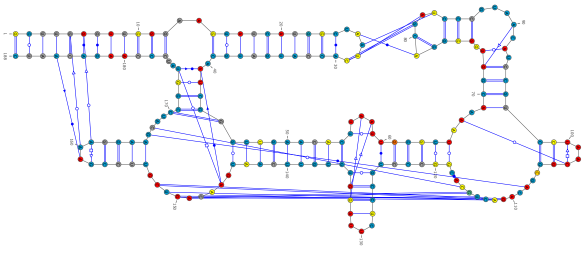

4 – NMIA-MaP$_{\text{IT}}$ (Cluster A ●)

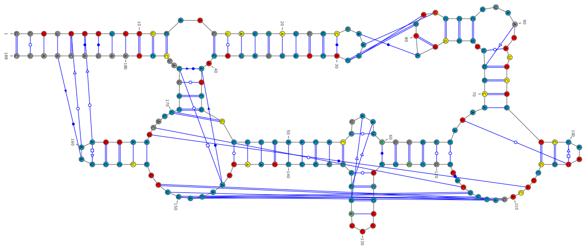

5 – 1M7-MaP$_{\text{IL}}^{\text{MG}}$-3D (Cluster C ●)

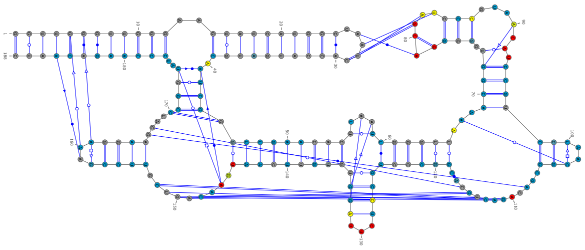

6 – NMIA-CE$^{\text{MG}}$ (Cluster E ●)

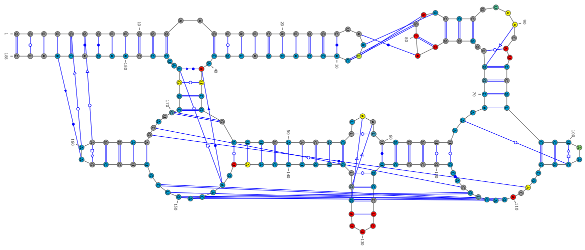

7 – NAI-CE$^{\text{MG}}$ (Cluster G ●)

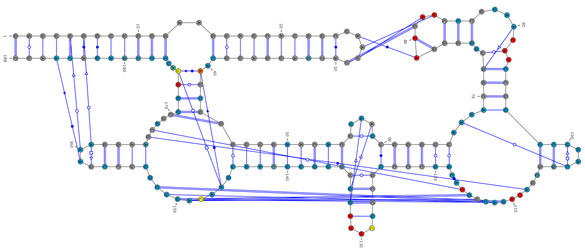

8 – BzCN-CE$^{\text{MG}}$ (Cluster H ●)

**Figure S2:** Projections of reactivities onto native structure (conditions 1-8 of 16).

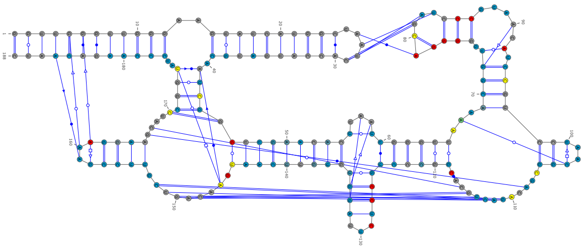

9 – 1M7-CE$^{MG}$ (Cluster E ●)

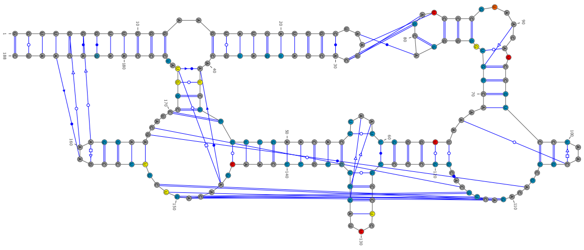

10 – CMCT-CE$^{MG}$ (Cluster G ●)

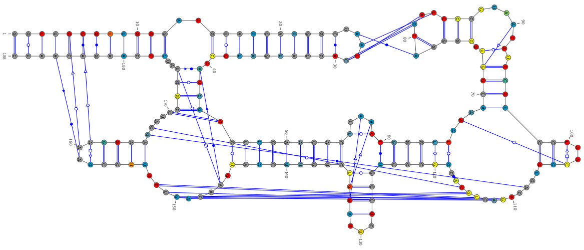

11 – 1M7-MAP$_{IL}$-3D (Cluster D ●)

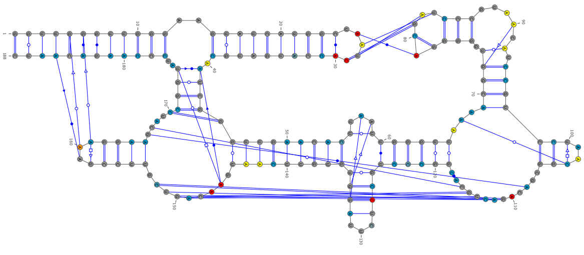

12 – DMS-CE$^{MG}$ (Cluster F ●)

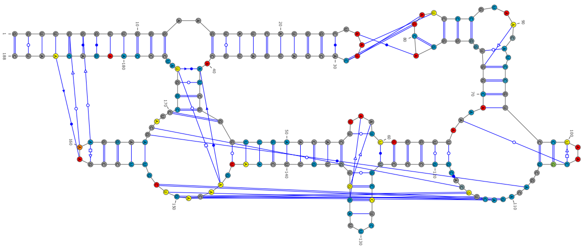

13 – NMIA-CE (Cluster B ●)

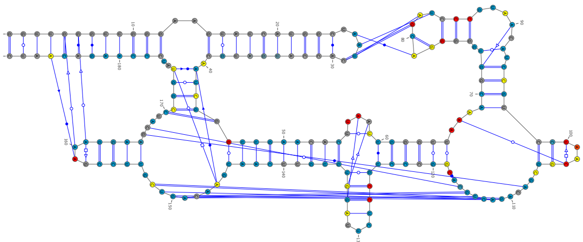

14 – 1M7-CE (Cluster B ●)

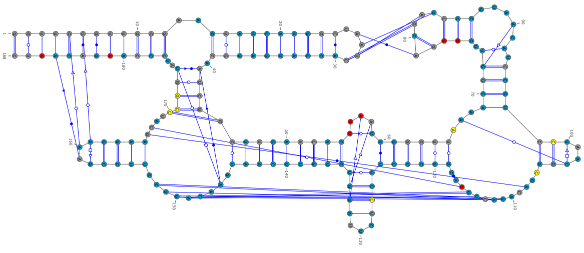

15 – BzCN-CE (Cluster B ●)

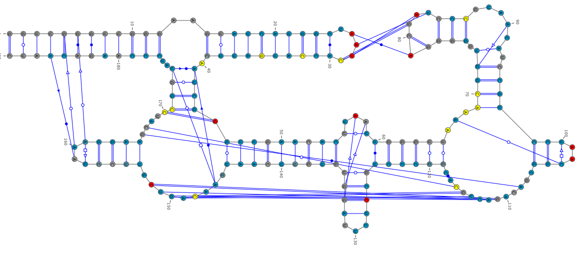

16 – NAI-CE (Cluster B ●)

**Figure S3:** Projections of reactivities onto native structure (conditions 9-16 of 16).

#### 3.2 Correlation analysis of raw reactivities profiles

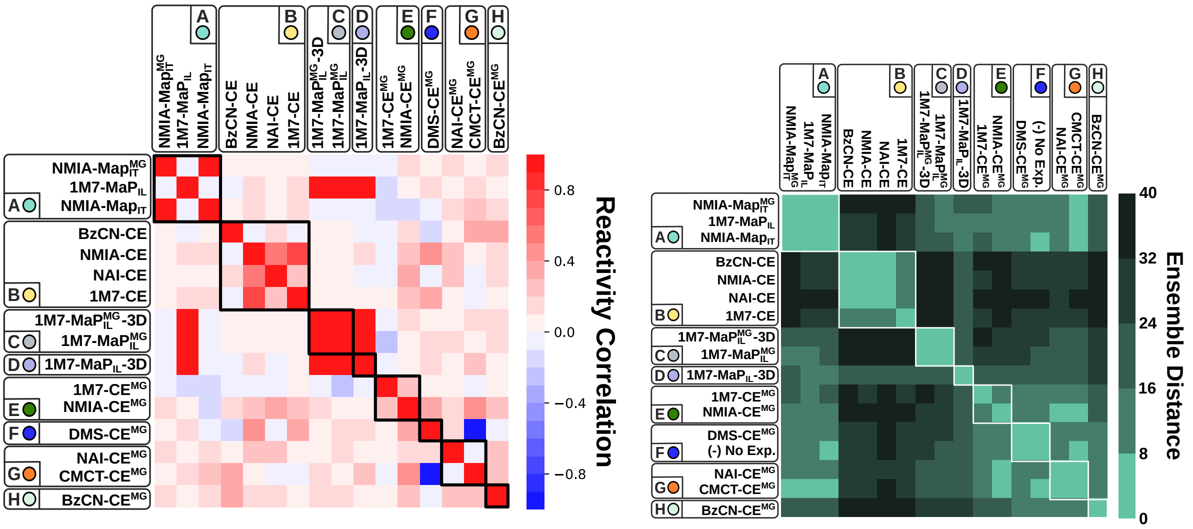

**Figure S4:** Correlation of reactivity profiles (left) and induced (pseudo-)ensemble distance (right). Conditions clustered in our analysis due to their impact of the pseudo-Boltzmann ensemble (black boxes, right) do not necessarily correlate very well in term of raw reactivity (white boxes, left).

We compared raw reactivity data produced across various conditions, in order to test their overall compatibility, investigate their main determinants and compare their impacts on predictions.

**Correlation metrics and rationale.** To that purpose, we consider the Pearson product-moment **correlation coefficients** $C_{r,r'}$ of two reactivity profile $r$ and $r'$, defined as

$$C_{r,r'} = \frac{\sum_{i=1}^{n} (r_i - \mu_r)(r'_i - \mu_{r'})}{\sqrt{\sum_{i=1}^{n} (r_i - \mu_r)^2} \sqrt{\sum_{i=1}^{n} (r'_i - \mu_{r'})^2}},$$

where $\mu_r$ and $\mu_{r'}$ denote the mean reactivity within $r$ and $r'$ respectively.

The choice of the **correlation coefficient as a measurement of pairwise similarity** for probing profiles is motivated by differences in the distributions of reactivities across experimental conditions and protocols. Indeed, $C_{r,r'}$ remains unchanged by the application of any rescaling, through an affine transformation with positive slope, to either $r$, $r'$ (or both). Moreover, this metrics does not scale with the number of positions. Thus, we can restrict the sums and mean values to positions for which reactivities are available in both experiments, and still compare the coefficient with another pair of reactivity profiles. This property is important in order to tolerate **missing values** introduced by experimental artifacts and reagent properties, while still being able to compare the coefficients.

**Results.** We computed the correlation coefficient for all pairs of reactivity profiles, and report the results in the left matrix of Figure 4. For the sake of comparison, we also remind in this Figure the Ensemble Distance matrix, indicating the difference in impact on the computational predictions of each pair of conditions.

We first observe that almost all of the pairs of reactivity profiles show a positive correlation, as expected from their observation of a common structure. A notable exception is 1M7-CE$^{MG}$, having average negative correlation with all its peers (except for NMIA-CE$^{MG}$). Its outlier status is confirmed by its large ensemble distance to other conditions, yet does not prevent it to achieve a respectable MCC of 70%. Finally, the perfectly negative correlation between CMCT-CE$^{MG}$ and DMS-CE$^{MG}$ can be dismissed as a computational artifact, since CMCT and DMS react with different nucleotides, and the correlation cannot be computed.

Moreover, we observe that conditions clustered from the Ensemble Distance metrics (due to their impact of the pseudo-Boltzmann ensemble) do not always correlate in term of raw reactivity, with very poor correlations occurring within clustered conditions. This essentially reflects the fact that similar profiles may induce very similar pseudo-Boltzmann ensembles, *ie* different premises sometimes imply the same conclusions.

Conversely, one observes very few strong correlations outside of clusters. In fact, the only two exceptions are 1M7-MAP$_{IL}$ and 1M7-MAP$_{IL}$-3D which, despite belonging to different clusters and achieving different qualities of prediction, correlate almost perfectly with each other, and with other reactivity profiles produced using 1M7/SHAPEMap. Other natural candidates for explaining the reactivity correlations (Map vs CE, +Mg vs -Mg…) do not appear to be supported by this analysis.

Overall, this analysis confirms the consistency of the produced probing profiles (positive correlation), but also suggests a very delicate effect of reactivity data on the downstream predictions, with very different profiles leading to similar predictions (1M7-MAP$_{IL}$), and similar profiles inducing very different ones (1M7-MAP$_{IL}$ vs 1M7-MAP$_{IL}$-3D).

#### 3.3 Dominant conformations within main clusters

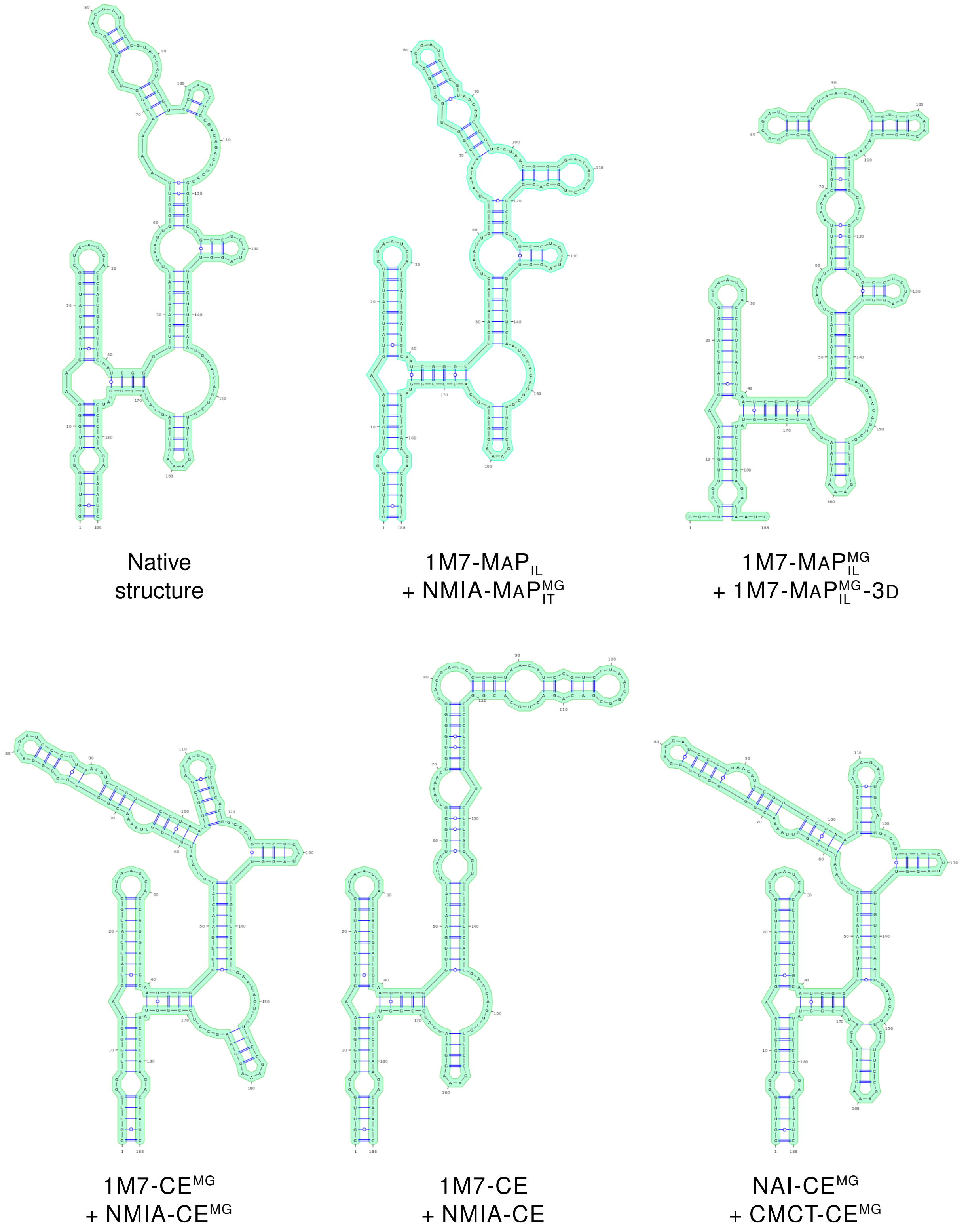

**Figure S5:** Predicted structures with conditions belonging to the same cluster

##### 3.4 Prediction accuracy for pairs of conditions

| | Mono | **1** | **2** | **3** | **4** | **5** | **6** | **7** | **8** | **9** | **10** | **11** | **12** | **13** | **14** | **15** | **16** |
|---|---|---|---|---|---|---|---|---|---|---|---|---|---|---|---|---|---|
| **1** – 1M7-MAP$^{MG}_{IL}$ | 0.85 | 0.85 | 0.84 | 0.8 | 0.82 | 0.82 | 0.82 | 0.75 | 0.79 | 0.78 | 0.79 | 0.79 | 0.85 | 0.61 | 0.79 | 0.79 | 0.85 |
| **2** – 1M7-MAP$_{IL}$ | 0.84 | | 0.84 | 0.82 | 0.82 | 0.7 | 0.83 | 0.72 | 0.82 | 0.85 | 0.83 | 0.84 | 0.84 | 0.84 | 0.6 | 0.84 | 0.6 |
| **3** – NMIA-MAP$^{MG}_{IT}$ | 0.81 | | | 0.81 | 0.8 | 0.74 | 0.82 | 0.82 | 0.81 | 0.81 | 0.79 | 0.8 | 0.8 | 0.61 | 0.81 | 0.81 | 0.6 |
| **4** – NMIA-MAP$_{IT}$ | 0.8 | | | | 0.8 | 0.74 | 0.73 | 0.81 | 0.78 | 0.7 | 0.79 | 0.63 | 0.72 | 0.61 | 0.7 | 0.7 | 0.59 |
| **5** – 1M7-MAP$^{MG}_{IL}$-3D | 0.74 | | | | | 0.74 | 0.74 | 0.7 | 0.74 | 0.74 | 0.7 | 0.63 | 0.74 | 0.74 | 0.74 | 0.74 | 0.74 |
| **6** – NMIA-CE$^{MG}$ | 0.73 | | | | | | 0.73 | 0.73 | 0.73 | 0.73 | 0.73 | 0.61 | 0.73 | 0.61 | 0.73 | 0.73 | 0.6 |
| **7** – NAI-CE$^{MG}$ | 0.73 | | | | | | | 0.73 | 0.72 | 0.7 | 0.73 | 0.64 | 0.68 | 0.61 | 0.73 | 0.73 | 0.59 |
| **8** – BzCN-CE$^{MG}$ | 0.71 | | | | | | | | 0.71 | 0.65 | 0.73 | 0.63 | 0.72 | 0.6 | 0.61 | 0.6 | 0.59 |
| **9** – 1M7-CE$^{MG}$ | 0.7 | | | | | | | | | 0.7 | 0.7 | 0.7 | 0.7 | 0.61 | 0.61 | 0.6 | 0.6 |
| **10** – CMCT-CE$^{MG}$ | 0.7 | | | | | | | | | | 0.7 | 0.63 | 0.74 | 0.61 | 0.6 | 0.6 | 0.6 |
| **11** – 1M7-MAP$_{IL}$-3D | 0.64 | | | | | | | | | | | 0.64 | 0.62 | 0.62 | 0.65 | 0.65 | 0.59 |
| **12** – DMS-CE$^{MG}$ | 0.63 | | | | | | | | | | | | 0.63 | 0.61 | 0.6 | 0.6 | 0.6 |
| **13** – NMIA-CE | 0.61 | | | | | | | | | | | | | 0.61 | 0.62 | 0.6 | 0.6 |
| **14** – 1M7-CE | 0.6 | | | | | | | | | | | | | | 0.6 | 0.61 | 0.59 |
| **15** – BzCN-CE | 0.6 | | | | | | | | | | | | | | | 0.6 | 0.6 |
| **16** – NAI-CE | 0.6 | | | | | | | | | | | | | | | | 0.6 |

**Table S6:** MCC of structures prediction with `IPANEMAP` from any pair of conditions

.

##### 3.5 Histograms of prediction accuracy informed by single-point mutants reactivities (Mutate-and-Map protocol)

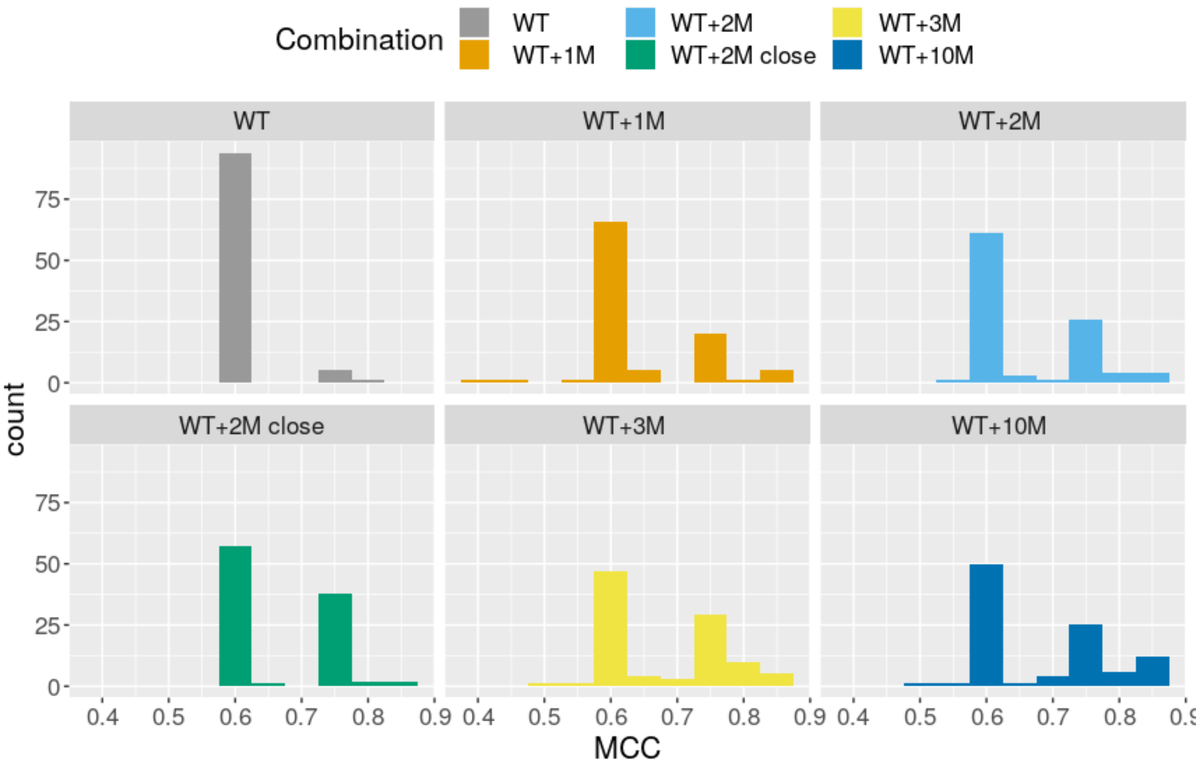

**Figure S6:** Distribution of MCC values over 100 runs from Mutate-and-Map data

#### References

[1] Cordero, P., Kladwang, W., Vanlang, C. C., and Das, R. (2012) Quantitative Dimethyl Sulfate Mapping for Automated RNA Secondary Structure Inference. *Biochemistry (Mosc )*, **51**(36), 7037–7039.

[2] Hajdin, C. E., Bellaousov, S., Huggins, W., Leonard, C. W., Mathews, D. H., and Weeks, K. M. (2013) Accurate SHAPE-directed RNA secondary structure modeling, including pseudoknots. *Proc Natl Acad Sci U S A,* **110**(14), 5498–5503.
